# Teasing apart the host-related, nutrient-related and temperature-related effects shaping the phenology and microbiome of the tropical seagrass *Halophila stipulacea*

**DOI:** 10.1101/2021.08.21.457214

**Authors:** Amir Szitenberg, Pedro Beca-Carretero, Tomás Azcárate-García, Timur Yergaliyev, Rivka Alexander-Shani, Gidon Winters

## Abstract

**Background:** *Halophila stipulacea* seagrass meadows are an ecologically important and threatened component of the ecosystem in the Gulf of Aqaba. Recent studies have demonstrated correlated geographic patterns for leaf endophytic community composition and leaf morphology, also coinciding with different levels of water turbidity and nutrient concentrations. Based on these observations, workers have suggested an environmental microbial fingerprint, which may reflect various environmental stress factors seagrasses have experienced, and may add a holobiont level of plasticity to seagrasses, assisting their acclimation to changing environments and through range expansion. However, it is difficult to tease apart environmental effects from host-diversity dependent effects, which have covaried in field studies, although this is required in order to establish that differences in microbial community compositions among sites are driven by environmental conditions rather than by features governed by the host.

**Results:** In this study we carried out a mesocosm experiment, in which we studied the effects of warming and nutrient stress on the composition of epiphytic bacterial communities and on some phenological traits. We studied *H. stipulacea* collected from two different meadows in the Gulf of Aqaba, representing differences in the host and the environment alike. We found that the source site from which seagrasses were collected was the major factor governing seagrass phenology, although heat increased shoot mortality and nutrient loading delayed new shoot emergence. Bacterial diversity, however, mostly depended on the environmental conditions. The most prominent pattern was the increase in Rhodobacteraceae under nutrient stress without heat stress, along with an increase in Microtrichaceae. Together, the two taxa have the potential to maintain nitrate reduction followed by an anammox process, which can together buffer the increase in nutrient concentrations across the leaf surface.

**Conclusions:** Our results thus corroborate the existence of environmental microbial fingerprints, which are independent from the host diversity, and support the notion of a holobiont level plasticity, both important to understand and monitor *H. stipulacea* ecology under the changing climate.

## Introduction

*Halophila stipulacea* (Forsk) Ascherson is a small tropical seagrass, dominant in the Gulf of Aqaba [1–3], the northernmost edge of its natural range [4, 5]. Here *H. stipulacea* forms large discontinuous meadows along a wide range of depths [1-50m; 2, 3, 6, 7]. Ecosystem services and functions associated with *H. stipulacea* meadows in this region are numerous. *H. stipulacea* are attributed with high primary productivity, enriching the water with oxygen. They sequester blue carbon, partially mitigating ocean acidification for neighbouring reefs [8]. They also reduce pathogen loads in seawater [9] and uptake nutrients, improving water clarity for neighbouring ecosystems [reviewed in 10]. Importantly, they provide major nursery grounds for fish, crustaceans, gastropods and bivalves [1]. However, these meadows are severely threatened by anthropogenic pressures. Seawater in the Gulf of Aqaba has been warming 50% faster than the global mean coastal sea surface temperature trend of 0.17 ± 0.11°C per decade [11, 12]. In addition, they are faced with intensive coastal development (e.g, the Saraya laguna in Aqaba, and the Splash Park in Eilat). These threats to seagrasses and the ecological functions they provide are made worse by the long water residence time estimates in the Gulf [3-8 years; 13] caused by its semi-enclosed basin shape, and the effects of eutrophication are intensified several-fold [reviewed by 2].

With the ongoing decline of seagrasses worldwide [14] alongside the relatively slow responding community based indicators in most seagrass monitoring efforts [15–17], there is a growing need for fast and responsive indicators to changes in the ecological status of seagrasses [18]. In a study of three Gulf-of-Aqaba *H. stipulacea* meadows growing at a shared depth, Mejia et al. [19] was able to distinguish bacterial “environmental fingerprints” for the different sites, from a “core microbiome”, which was shared among all sites. Bacterial differences among sites were attributed mostly to Rhodobacteraceae, which dominated particularly at the southernmost meadow, the site with highest nitrogen and phosphate concentration measurements, but with lowest turbidity. Additionally, plants from this meadow had the smallest leaf area, among the studied meadows. The existence of “environmental fingerprints” alongside a core microbiome was also corroborated by Rotini et al. [7], who compared the microbiomes in *H. stipulacea* meadows along a depth gradient of 4-28m. The epiphytic leaf microbial communities of these meadows were found to be dominated by Gammaproteobacteria and Bacteroidetes in the light limiting conditions of the deeper sites, while Cyanobacteria and Rhodobacteraceae thrived in conditions of high light availability and hydrodynamics, in shallower sites. Based on the “environmental fingerprint” hypothesis, bacterial community shifts, and particularly Rhodobacteraceae relative abundance increase, may provide early indications to nutrient exposure [7, 19, 20], and possibly act as a buffer, protecting the seagrass host from excess nutrients through bacterial metabolism. However, field studies cannot tease apart the effects of environmental factors from those of possible differences in host diversity among seagrass populations.

Outside of its natural range, *H. stipulacea* is a successful lessepsian migrant in the eastern Mediterranean Sea [21–25] and an invasive species in the Caribbean Sea [26, 27]. The ability to migrate into new habitats could be related not only to its extensive morphological, biochemical and physiological plasticity [reviewed by 2, 6, 7, 28–30], but perhaps also to its bacterial “environmental fingerprints”. Like many other organisms, seagrasses were suggested to establish symbiotic relationships with their associated microbial communities to form a functional unit (the “holobiont”) that reacts as a whole to environmental changes [31–33]. It has been suggested that the epiphytic bacterial community of the seagrass leaves, the focus of this study, benefit from the organic carbon enriched microhabitat on the leaves [34] and comprises aerobic organotrophic bacterial species able to utilize the secreted polymers [7, 34]. Diazotrophic bacteria such as cyanobacteria, which enhance nitrogen availability [35–38], may also be a part of this community, depending on nutrient concentrations in the water [39]. Alternatively, leaf cyanobacterial biofilms may reduce light availability [40].

In this study we aimed to understand whether the bacterial “environmental fingerprint” in *H. stipulacea* existed independently from host related diversity, which may exist among meadows in the field, in order to establish the utility of bacterial shifts as early warning signs for environmental stress. This would also provide evidence for holobiont level plasticity, supporting the rapid range extension of *H. stipulacea*. We also tested the interaction between phenotypic responses of *H. stipulacea*, including shoot production and mortality, with microbial community shifts, under temperature and nutrient stress. We carried out a mesocosm experiment in which seagrass phenology and leaf epiphyte community compositions were the dependent variables, while the source seagrass population, water temperature and nutrient concentrations were the independent variables. We simulated shallow meadow conditions of 8-10 m depth, in which light was not a limiting factor. We hypothesised that both the environmental conditions and the source seagrass population would partially explain the variance in the dependent variables, and that the interaction of the independent factors would affect the seagrass performance and the microbiome differently than each factor separately.

## Results

To study the effects of heat, nutrients and the source *H. stipulacea* population on the phenology and epiphytic microbiome of *H. stipulacea* in a controlled environment, we carried out a mesocosm experiment. We controlled for the source population by including seagrasses from two sites, Tur-Yam Beach (TY) and South Beach (SB), both located in the northern Gulf of Aqaba (Eilat, Israel). TY is offshore a small and active marina, close to a rarely used crude oil terminal, whereas SB is removed from any obvious anthropogenic pressures. While turbidity in the TY site is slightly higher than in the SB site, higher nutrient concentrations were measured in SB (further detailed in Mejia et al. 2016). To reduce the effect of microbial legacies, seagrasses were washed with freshwater and re-inoculated by exposure to natural seawater. The mesocosm aquaria system (Fig. 1) included four temperature baths, each containing five aquaria filled with artificial seawater. Following acclimation, under 27°C and no nutrient enrichment, the mesocosm conditions diverged to four different regimes, for the duration of the 40 days experimental phase: *i*) control water temperature with no nutrient enrichment (CTCN), *ii*) control water temperature with nutrient enrichment (CTN), *iii*) increased water temperatures (31°C) with no nutrient enrichment (TCN), and *iv*) the combination of the two stressors - increased water temperatures (31°C) with nutrient enrichment (TN). The experimental phase was followed by a recovery step in which baseline temperatures were set in all the baths and nutrient addition was stopped.

**Fig. 1:**
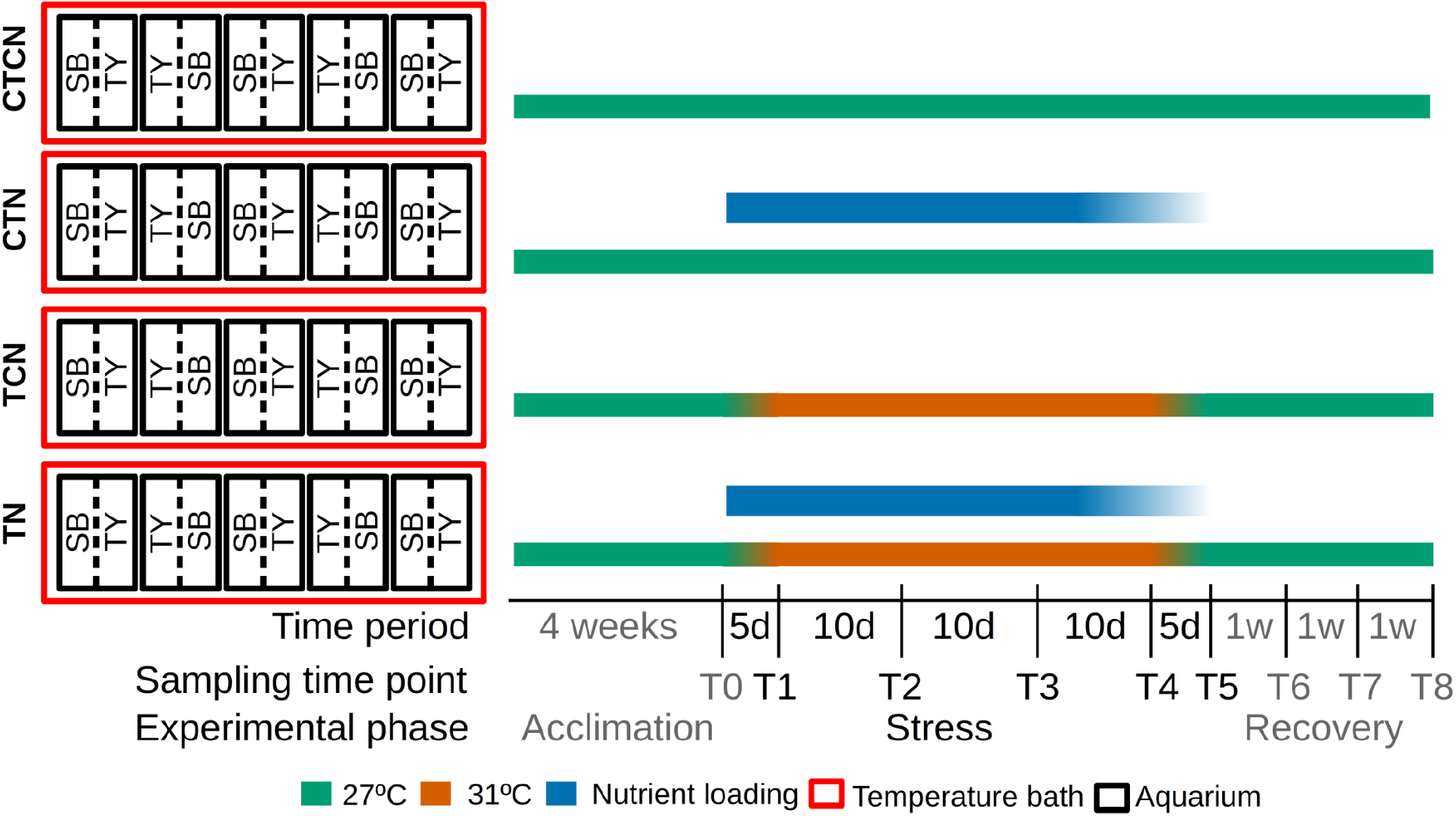
Mesocosm and experimental setup. Seagrasses collected from South Beach (SB) and Tur-Yam Beach (TY) were planted in two divisions (broken line) within aquaria (black rectangles). The aquaria were distributed among four temperature baths. The experimental timeline with sampling timepoints (T0 - T8), the elapsed time between sampling time points and the experimental phase they belong to are denoted at the bottom. Bath temperatures are denoted by green (27°C) and orange (31°C) bars. Active maintenance of high nutrient concentrations is denoted with blue bars. CTCN - control temperatures (27°C) and control nutrients (no loading). CTN - control temperatures with nutrient loading . TCN - heatwave (31°C) without nutrient loading. TN - heatwave with nutrient loading. Time point 0 & 8 had baseline conditions in all baths.

### Nutrient measurements, shoot formation and shoot mortality

The establishment of heatwave conditions (31°C) in TCN and TN aquaria during the stress phase of the experiment (T1 - T5) is demonstrated in Fig. 2A. NO_2_^−^ concentrations (Fig. 2B) in the non-enriched aquaria (CTCN, TCN) remained similar to the baseline values throughout the experiment (0.08 - 0.16 μM on average), and were lower than the enriched aquaria(CTN, TN) at T3 and T5 (0.28 - 0.63 μM on average; 0.026 < p-value < 0.03). Interestingly, following recovery, NO_2_^−^ concentrations returned to baseline, pre-treatment values in all aquaria except CTN, where they were elevated (0.32 μM on average; p-value < 0.003). A corresponding decrease in NO_3_^−^ concentrations was observed under the same treatment (Fig. 2C). NH_4_^+^ concentrations (Fig. 2D) seemed less affected by the enrichment treatment. By T3 only the CTN treatment aquaria had a clear elevated level of NH_4_^+^ (2.99 μM on average; p-value < 0.023) compared to NH_4_^+^ concentrations in non enriched aquaria (1.01 and 1.46 μM on average). By T5, only TN aquaria, exposed to thermal stress PO_4_^3−^ and enriched with nutrients, had such excess (1.99 μM on average). Following recovery, elevated NH_4_^+^ concentrations were observed in all aquaria (3.21-4.4 μM on average). PO_4_^3−^ concentrations (Fig. 2E) were similar in all treatments at T0 (1.36-1.62 μM on average), diverging between the non-enriched and nutrient enriched treatments at time points T3 and T5 with values of 1.01-1.46 μM and 1.85 - 2.99 μM on average for non-enriched (CTCN, TCN) and nutrient enriched (CTN, TN) treatments, respectively (0.0187 < p-value < 0.068), as expected. PO_4_^3−^ concentrations at T8 were very high, albeit similar among treatments (3.21 - 4.40 μM on average), most likely due the the accumulation of PO_4_^3−^ in all the aquaria.

**Fig. 2:**
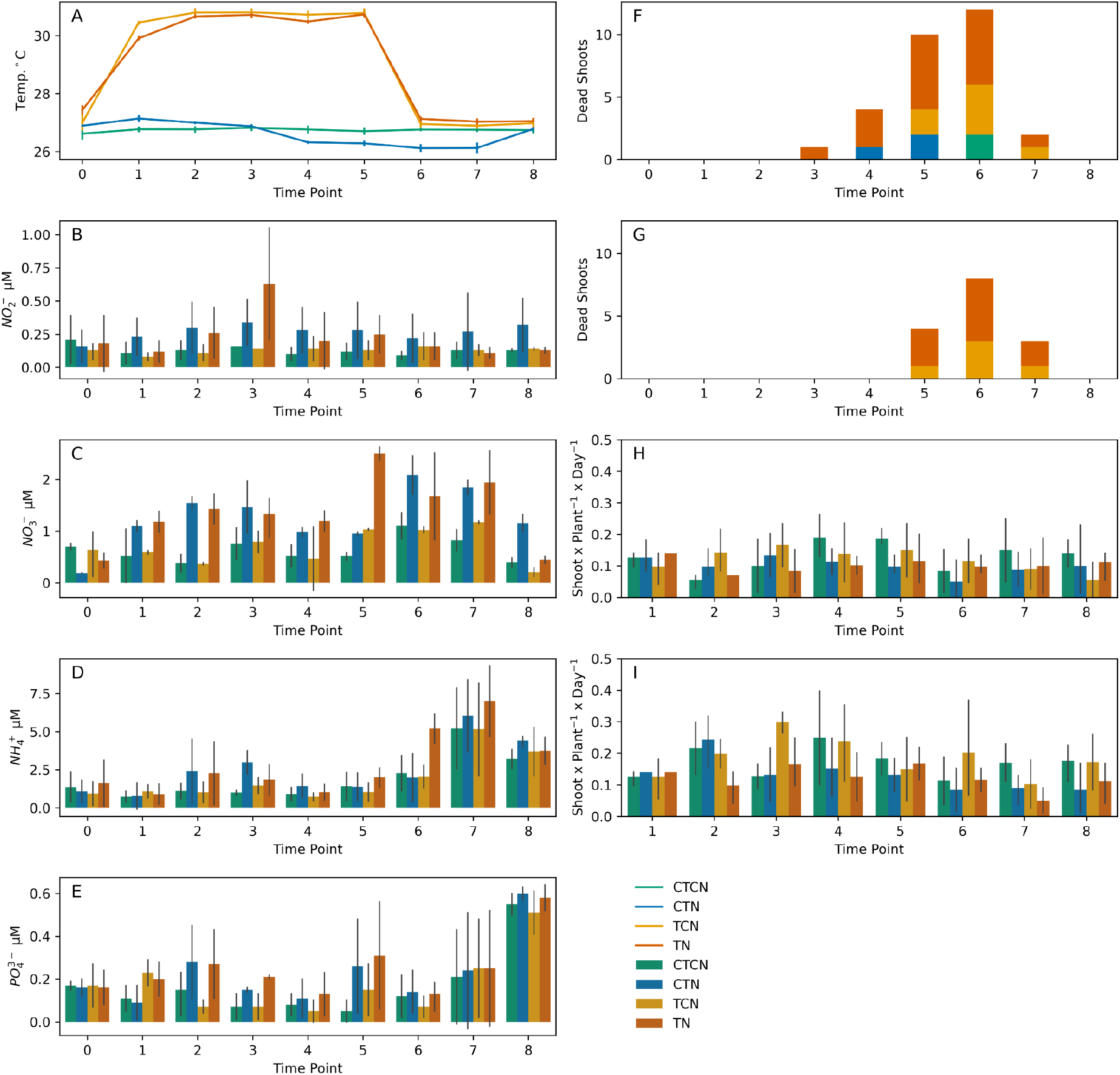
Changes over time in (A) daily average water temperatures (°C), concentrations (μM) of nitrite (B), nitrate (C) ammonium (D) and phosphate (E), shoot mortality in SB (F) and TY (G) plants, and per-plant, per-day mean shoot additions between time points in SB (H) and TY (I) plants. Nutrient concentrations represent measured concentrations, which could be perturbed by accumulation and biological activities, and not the amount of loaded nutrients. CTCN - control temperatures (27°C) and control nutrients (no loading). CTN - control temperatures with nutrient loading . TCN - heatwave (31°C) without nutrient loading. TN - heatwave with nutrient loading. Time point 0 & 8 had baseline temperatures, without nutrient enrichment, in all baths.

Shoot mortality appeared among TY plants only at T5, and only in heated aquaria (TCN, TN), in which a total of 15 dead shoots were counted (Fig. 2G). Shoot mortality in SB plants, however (Fig. 2F), was evident two weeks earlier at T3 with a total of 29 dead shoots across treatments and time points, including five from unheated aquaria (CTCN, CTN). Correspondingly, new shoot emergence (Fig. 2H & I) was significantly higher in TY plants than SB plants (F = 16, p-value = 8 x 10^−5^) and in non-enriched aquaria (CTCN, TCN) than in enriched aquaria (CTN,TN) with F = 4.6 and p-value = 0.003, in a type-3 factorial ANOVA. Time was a marginally significant factor (F = 2.3, p-value = 0.02). Therefore, according to phenology measurements, TY seagrass inherently performed better under thermal stress for both shoot death and emergence, temperature stress primarily promoted shoot death, more so in SB hosts more than in TY hosts, and nutrient loads suppressed the emergence of new shoots for both SB and TY hosts, down to a similar level.

### Sequencing results

To study the temporal dynamics of the leaf epiphytic microbiota, we carried out a metabarcoding experiment. Following the exclusion of organelle sequences and chimeric sequences, as well as amplicon sequence variants (ASVs) with a frequency lower than 30 across all samples, the analysis included 142 mesocosm epiphyte samples, 20 mesocosm water samples and 10 field epiphyte samples, with a median read-pair count of 14,501. Except for one of the retained mesocosm epiphyte samples, which had 5,851 reads, the filtered read count ranged from 8,312 to 33,256.

### Bacterial diversity

Samples were rarified to 5000 read-pairs following the guidance of a rarefaction curve, which confirmed that most rare taxa were represented (Fig. S1). Water samples had significantly lower Faith PD values than epiphyte samples (ANOVA p-value < 10^−56^; Fig. S2). The first principal coordinate in a principal coordinate analysis (PCoA; Fig. S2), explained 38.4% of the total variance and segregated the sample types, water vs. endophytes, to distinct clusters. Water samples were more divergent from one another than endophyte samples, and the main ASVs explaining the difference between the sample types belonged to the family Alteromonadaceae (ASV cc2916), the NS3a marine cluster (745c89) and the genus *Winogradskyella* (01f97d), which were more abundant in the water samples (Fig. S2).

To study the bacterial diversity within the endophytic communities, we excluded ASVs that were also found in the aquarium-water samples and regarded them as contamination. Faith PD values were similar among time points and between the source sites (SB and TY). Treatments were a significant factor (ANOVA p-value = 0.01), due to significant differences between the TN and CTN treatments (q-value = 0.005; Fig. 3), both of which were enriched at T3 and T5, differing only in the water temperatures applied. The decrease in alpha diversity under CTN corresponded with the increase in NO_2_^−^ under the same treatment (Fig. 2B).

**Fig. 3.**
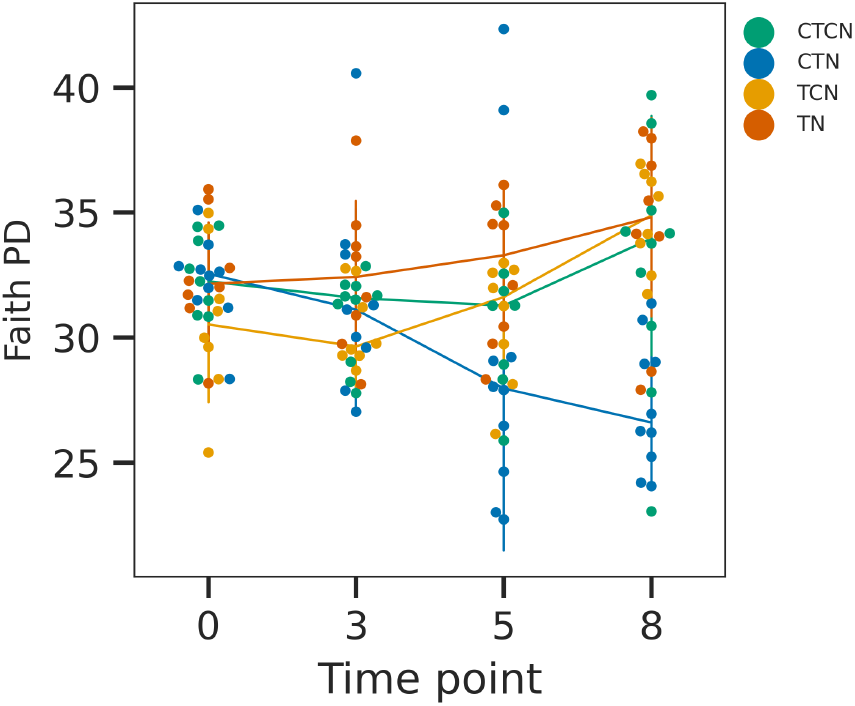
Faith PD temporal dynamics. The four treatments (CTCN - control temperatures (27°C) and control nutrients (no nutrient enrichment). CTN - control temperatures with nutrient enrichment. TCN - heatwave (31°C) without nutrient enrichment. TN - heatwave with nutrient enrichment) are color coded. Line-plots represent the median Faith PD values and the bars denote standard deviations. Time point T0 and T8 had baseline conditions in all baths.

The unweighted UniFrac PCoA analysis (Fig. 4) revealed an increase in the divergence among samples from different treatments, following T0. As time progressed, the second axis (8%-9% of the total variance) segregated the control samples (CTCN) from the rest of the treatments, and axis 1 (12%-13%) separated CTN from the other treatments. The first and second axis in the Weighted UniFrac analysis (Fig. S3) explained a larger cumulative portion of the total variance (25%-30%) with similar patterns but shorter distances among the treatments. According to the redundancy analysis ANOVA, a model accounting for time, temperature, source population and the concentrations of NH_4_^+^, NO_2_^−^, NO_3_^−^ and PO_4_^3−^, accounted for 21% of the total variance (p-value = 0.001), with small but significant effects of time (3.3%, q-value=0.003), temperature (2.9%, q-value=0.003), PO_4_^3−^ concentration (2%, q-value=0.007), NO_2_^−^ (1.2%, q-value = 0.04), NO_3_^−^ (1.8%, q-value = 0.008) and NH_4_^+^ (1.2%, q-value=0.038). The source site had a borderline significant effect (q-value=0.09) explaining 1% of the variance. The remaining 7.5% of the variance explained by the model can be attributed to interactions among these factors.

**Fig. 4:**
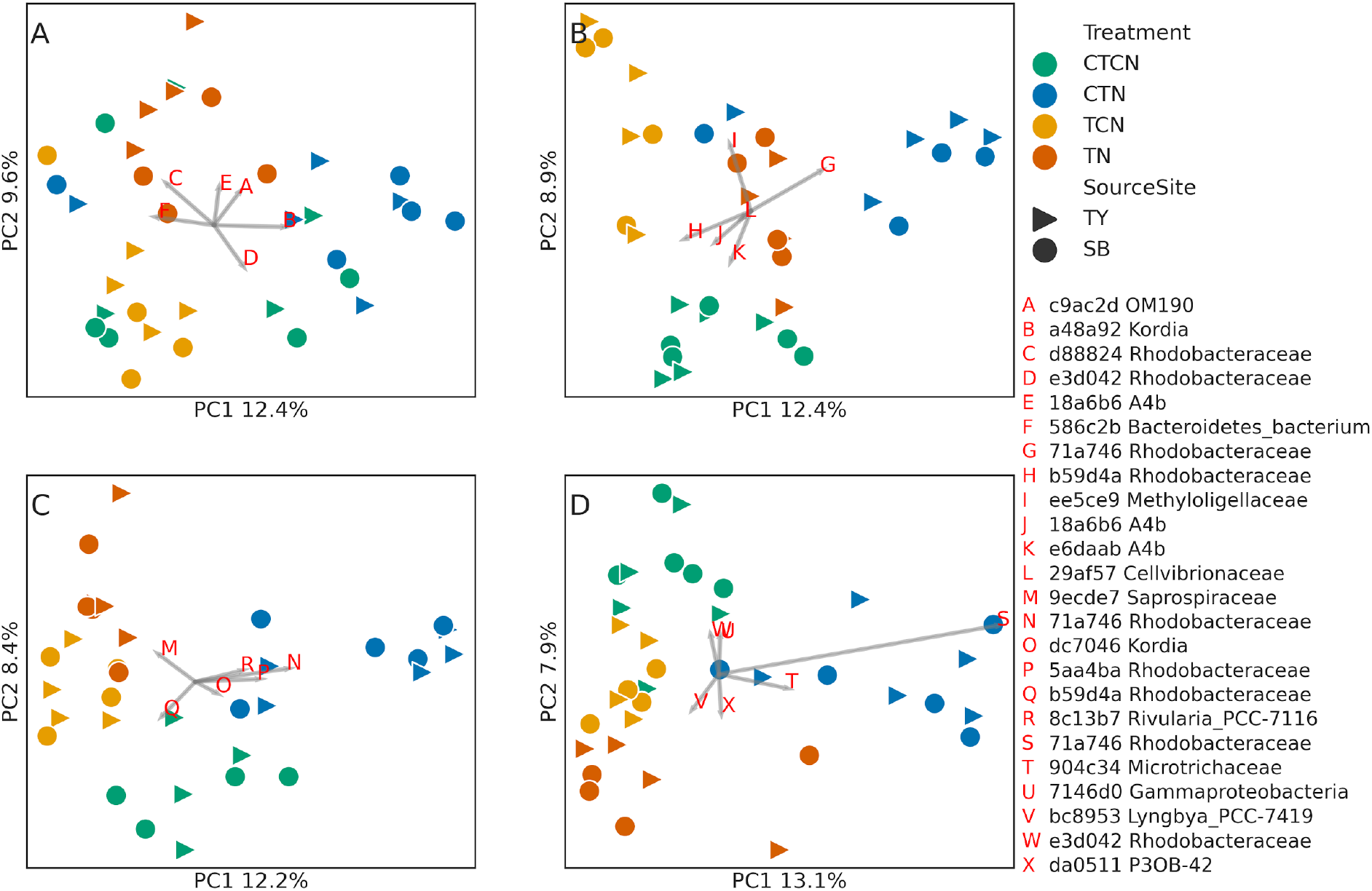
Diverging community compositions among treatments. Unweighted UniFrac distance based principal coordinate analyses (PCoA) of endophyte samples from T0 (A), T3 (B), T5 (C) and T8 (D). CTCN - control temperatures (27°C) and control nutrients (no enrichment). CTN - control temperatures (27°C) with nutrient enrichment. TCN - heatwave (31°C) without nutrient enrichment. TN - heatwave with nutrient enrichment. Time points T0 and T8 had baseline temperatures and no active nutrient enrichment in all baths. The percent total variance accounted for by each coordinate is indicated on the corresponding axis. The most important ASVs, following the importance definition by Legendre and Legendre [41], are represented by BiPlot analyses (gray arrows) and their taxonomic identifications are noted in the legend.

To further understand the importance of the source site in determining the response of biofilm communities to the treatment, we carried out a factorial ANOVA test using UniFrac pairwise distances. We tested whether within-SB, within-TY, and between-sites pairwise distances were significantly different (Fig. 5). We took care to only include within-treatment and within-time point pairwise distances and to test whether the selected distances differed among the treatments.

**Fig. 5:**
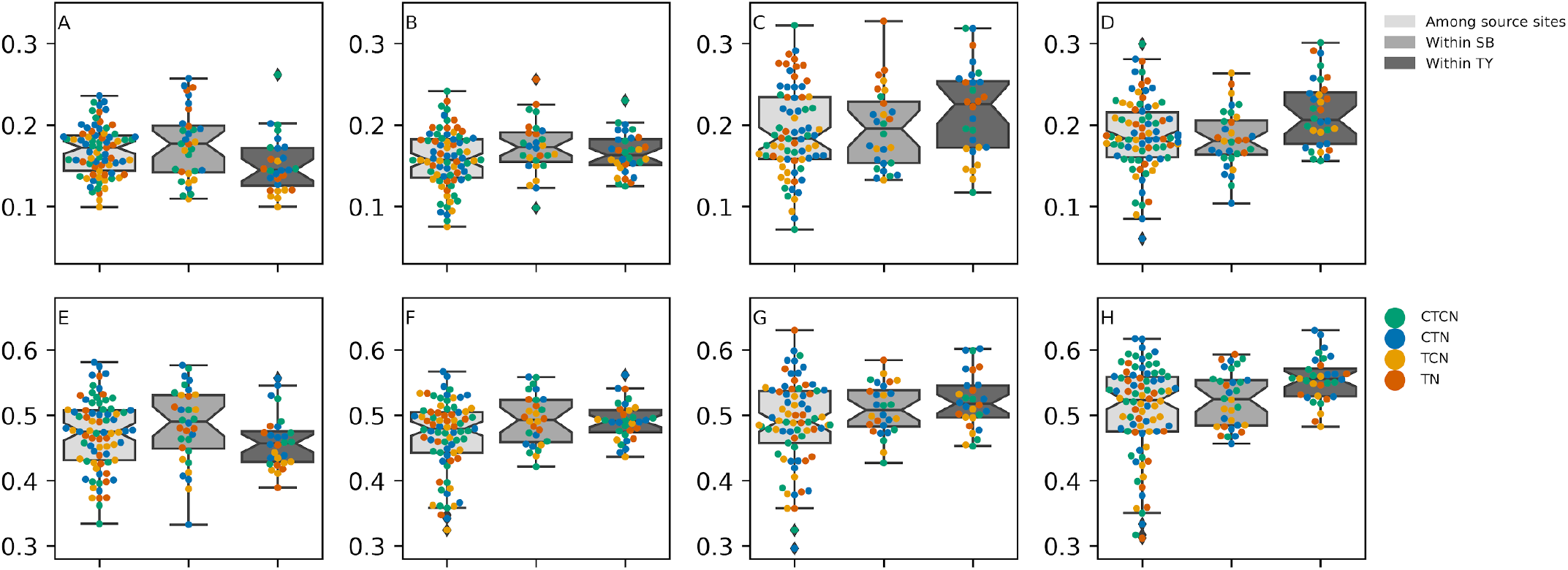
Dynamics of UniFrac pairwise distances within and among source sites along the experimental time frame. The distributions of among-site, within SB and within TY distances are presented as box plots (see legend), using weighted (A-D) and unweighted (E-H) distances. Time points 0 (A & E), 3 (B & F), 5 (C & G) and 8 (D & H) are presented separately. The swarm plots reflect the distribution of pairwise distances among the different treatments.

At T0, pairwise distances within TY were smaller than within SB or between source sites when considering weighted distances (p-value < 0.012) but not when using unweighted distances. In intermediate time points (Fig. 5C & G), particularly in T5, pairwise distance increased both within and between source sites, with a greater increase in the within-TY distances, (q-value = 0.047 and q-value = 0.024 for weighted and unweighted distances, respectively). Following recovery (Fig. 6 D & H), the within-TY distances increased further compared with the within SB distances (q-value = 0.005 for weighted distances) or the among-site distances (q-value = 0.005 and q-value = 5 x 10^−4^). It would therefore appear that the source site of the seagrass (the location from which the population was collected from) was one of the factors shaping the bacterial composition of the seagrass biofilm. Specifically, TY epiphytic communities have diverged from each other at higher rates than SB epiphytic communities. This result coincides with the increased mortality and reduced generation of shoots that was observed in SB plants compared with TY plants (Fig. 2F-I; mentioned above).

**Fig. 6:**
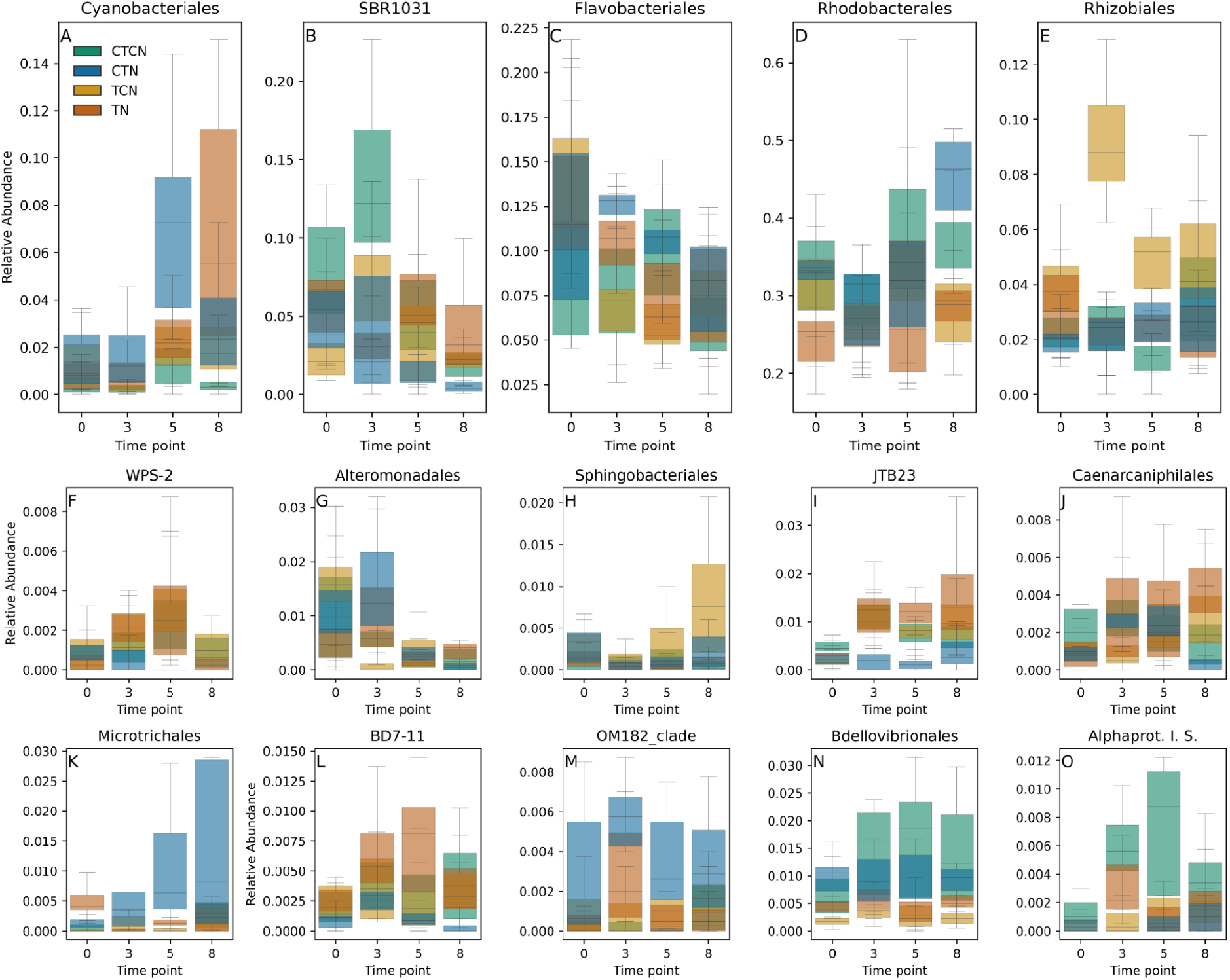
Order level dynamics of high abundance orders (A-E) and ANCOM orders (F-O). CTCN - control temperatures (27°C) and control nutrients (no enriching). CTN - control temperatures with nutrient enriching. TCN - heatwave (31°C) without nutrient enriching. TN - heatwave with nutrient enriching. Timepoints T0 and T8 had baseline temperatures and no active nutrient enrichment in all treatments.

### Order dynamics

Relative abundances of the high abundance orders were similar among treatments and time points (Fig. 6A), except for a few but important exceptions. The most abundant order (27% to 40%), Rhodobacterales (Alphaproteobacteria; Fig. 6C), had similar relative abundances in all treatments throughout the experiment, except for T8. By that time point, in which similar recovery conditions were already applied for three weeks in all the treatments, their relative abundance significantly increased (q-value < 0.009) under the legacy of control temperature conditions (27°C) and particularly under the legacy of control temperatures with high nutrients concentration (TCN), in correspondence with the increased NO_2_^−^ concentrations (Fig. 2).

Flavobacteriales (Fig. 6C, Bacteroidetes; Flavobacteriia) diverged among the treatments, with increased relative abundances under nutrient enriching (CTN & TN) at time point T3 (q-value = 0.025) and under control temperatures (CTCN & CTN) at time point 5 (q-value = 0.048), but the relative abundance in the different treatments converged under recovery conditions at T8. Mean relative abundances of Flavobacteriales ranged from 6% to 14% across time points and treatments (Fig. 6C). Similarly, Rhizobiales (Fig. 6E, Alphaproteobacteria) diverged among treatments at time points T3 and T5 (q-value < 0.02), to converge at T8, after three weeks of recovery conditions. However, Rhizobiales (Fig. 6E) consistently flourished only under heatwave conditions without nutrient loading (TCN). Their mean relative abundances ranged from 1.5% to 9% across time points and treatments. SBR1031-clade bacteria (Fig. 6B, Chloroflexi, Anaerolineae), with mean relative abundances of 2%-13%, had a higher relative abundance under control conditions (CTCN) at T3 (q-value < 0.03), but this increase did not persist, and this order was not among the eight most abundant orders following recovery. Cyanobacterales (Fig. 6A) emerged as a high relative-abundance order (2%-8%) at T5. They had higher relative-abundance under baseline temperatures with nutrient enriching (CTN) in T5 (q-value < 0.02).

ANCOM tests revealed additional differently abundant orders, with low relative abundances (Fig. 6F-O). Alphaproteobacteria incertae sedis (Fig. 6O, an artificial group of several alphaproteobacteria genera with unclear placement) were more abundant in the control samples (CTCN) than in any of the treatment samples. Alteromonadales (Fig. 6G; Alteromonadaceae in particular, Fig. S4I), decayed with time under all the experimental and control regimes. BD7-11 (phylum Planctomycetota; Fig. 6L) and Myxococcales (Deltaproteobacteria) followed a similar pattern to one another, where, unlike Microtrichales and Rhodobacterales, they were suppressed under CTN. Caenarcaniphilales (Fig. 6J) increased under stress compared to the T0 abundance in each treatment (CTN, TCN, TN), and flourished after recovery from heatwave and nutrient loading conditions (TN), JTB23 (Gammaproteobacteria; Fig. 6I), which increased under both heatwave treatments (TCN, TN), Microtrichales (Fig. 6K; particularly Microtrichaceae, Fig. S4G), which flourished under nutrient loading with baseline temperatures (CTN), similarly to Rhodobacterales, the OM182 clade (Gammaproteobacteria; Fig. 6M), which increased under nutrient loading conditions (CTN, TN) and Sphingobacteriales (Fig. 6H; particularly the NS11-12 marine group, Fig. S4F), which flourished during recovery from heatwave and baseline nutrients conditions (TCN). Additional orders differed among treatments but were consistent with their relative abundance at T0, prior to the stress phase of the experiment.

A family level ANCOM analysis (Fig. S4) largely mirrored the order level results, with the addition of the following families that did not belong to the above mentioned orders. Microscillaceae (Fig. S4C; order Cytophagales), which developed with time only under control conditions, Phormidiaceae (Fig. S4D; order Oscillatoriales), which developed best under nutrient loading conditions (CTN, TN) and flourished following recovery from the combined stress TN conditions and Rubinisphaeraceae (Fig. S4A; Planctomycetales), that increased under control temperature combined with nutrient enrichment conditions (CTN) and persisted even during the recovery.

### Dynamics of key ASVs

The increase in the relative abundance of ASV 71a746 (Rhodobacterales), which is highly explanatory of the total variance (Fig. 4) seems to be responsible for the observed increase in the relative abundance of Rhodobacterales, which corresponds with the decrease in Faith PD observed under CTN (Fig. 3). ASV 71a746 is most closely related to AM691091, an extremophile from the East German Creek system in Canada [42] and to several *Roseobacter* sequences (Fig. S5), a genus containing temperate and polar species [43], and was first described in seaweed [44]. Some *Roseobacter* reduce NO_3_^−^ to NO_2_^−^ [44]. The Microtrichales ASV 904c34 consistently developed only under the CTN conditions as well. Microtrichaceae can oxidize NH_4_^+^ with NO_2_^−^ via the anammox system [45]. Methyloligellaceae ee5ce9 (Rhizobiales) are methanotrophs [46], which flourished most under TCN. Dynamic cyanobacterial ASVs included Rivularia 8c13b7, particularly successful under CTN, which appears to be related to extremophilic cyanobacteria from alkaline, saline and thermal environments (Fig. S6). It was the most abundant cyanobacterial ASV, but it was displaced by T8. *Pseudanabaena* 2f6e75, another cyanobacterium, is closely related to an isolate from the sponge *Axinella damicornis* [KY744814; 47]. It emerged at T8 in all treatments, including CTN. Lyngbya bc8953, which is closely related to cyanobacterial isolates from the intestinal tract of herbivorous marine fish (HM630185) and black band disease coral tissue [DQ446127; 48], also emerged at T8, under TN. Saprospiraceae a2fde72, which is closely related to isolates from the surface of macroalgae (Fig. S7; DQ269042) prevailed under all treatments, except for TN. Under the TN treatment, this ASV was displaced by another Saprospiraceae ASV (9ecde7), but reemerged following recovery. Interestingly, ASV 9ecde7 is related to isolates from a Guerrero Negro hypersaline microbial mat [49]. In the SBR1031 group, which was most successful under control conditions, ASV A4b f35075 appeared to be an exception. It emerged late under TN. The gammaproteobacterium *Granulosicoccus* ASV 9d7587, flourished only under CTN, similarly to Microtrichales and Rhodobacterales. This ASV emerged as important in Fig. S3. *Granulosicoccus* was first identified in Antarctica, has low optimal temperatures, and can reduce nitrate [50]. The temporal dynamics of key ASVs and taxa are summarized in Fig. S8.

### Core ASVs

To evaluate the relevance of the results to natural communities we established the baseline core epiphytic microbiota including epiphytic microbiota from unmanipulated field samples collected specifically for this purpose, and all the samples of at least one of the treatments. All the recovered core ASVs existed in all the treatments, except for four TY core ASVs that were not observed under TN. In total, 107 core ASVs were recovered, 60 in both source sites (Fig. S9), 35 in TY only (Fig. S10) and 12 in SB only (Fig. S11). Site specific ASVs occurred in the other site as well, but did not meet the core ASV criteria. Alphaproteobacteria included 48 core ASVs, belonging to Rhodobacterales (37), Micavibrionales, Rhizobiales, Sphingomonadales (two each), Caulobacterales, Kiloniellales, Rhodospirillales, Rickettsiales and unidentified (one each). Bacteroidia included 25 core ASVs, belonging to Flavobacteriales (11), Chitinophagales (8) and Cytophagales (6). Gammaproteobacteria included 21 core ASVs, belonging to Cellvibrionales (7), Alteromonadales (5), unidentified (3), Gammaproteobacteria_Incertae_Sedis (2), Arenicellales, Burkholderiales, SZB50 and Steroidobacterales (one each). Rhodobacterales, Flavobacteriales, Rhizobiales, Sphingomonadales and Alteromonadales were all identified as orders responding to the experimental treatments (Fig. 6). Four ASVs recovered as “important” by the BiPlot analysis (Fig. 4 and S3) were also found among the core ASVs, shared among the source sites, including Koridia dc7046 (Flavobacteriales), Methyloligellaceae ee5ce9 (Rhizobiales), Rhodobacteraceae 71a746 and Rhodobacteraceae b59d4a (Rhodobacterales).

## Discussion

Studies of *H. stipulacea* meadows in the Gulf of Aqaba have revealed associations between the epiphytic above-ground microbiota composition and water nutrient concentrations [7, 19, 20]. Based on the observed associations, these studies have suggested that microbiota function within the seagrass holobiont and have pointed out their potential use as an ecological indicator of exposure to environmental stress by the seagrass host [20]. While *in-situ* studies might provide a better representation of realistic conditions to which seagrasses and their microbiota are exposed, it is often difficult to tease apart the relative importance of covarying factors, such as heat waves, nutrient loading and the host diversity. In addition, in field experiments, there is the possibility that legacy environmental conditions, rather than those measured during the experiment, are responsible for the observed bacterial community compositions [51]. This may lead to misinterpretation of the key factors, and of the meaning of bacterial dynamics as bioindicators of environmental stress. In this mesocosm experiment we set out to control for the temperature, nutrient concentrations and source meadow, to study their respective importance and to test the existence of an “environmental fingerprint” properties previous workers attributed to seagrass microbiota [7, 19, 20]. A fundamental requirement for such a marker is that bacteria that were indicated as key players in-situ are also represented in the experimental system at the end of the acclimation step (T0), so that the observed consequent dynamics will bear relevance to the natural meadows. This equivalence between the experimental system and previous in-situ results are best reflected by the core ASVs. The mesocosm study system has 107 core ASVs shared between the natural (in situ) samples and the experimental samples, including ASVs belonging to taxa which responded to the experiment or that were specifically shown to respond to the experiment. The largest cohorts of core ASVs belonged to alpha-proteobacteria (48 ASVs, mainly Rhodobacterales) and Bacteroidia (25 ASVs, mostly Flavobacteriales), both key groups of the aboveground *in situ H. stipulacea* microbiota [19]. As a case in point, the influence of the core ASV 71a746 on the alpha diversity in the mesocosm was paramount.

Two groups of dependent variables were quantified in this study, phenological and microbial. The two groups had fundamentally different governing factors. Phenological properties were largely dependent on the seagrass collection site, with overall better performance of the *H. stipulacea* from the TY. The two phenological traits, shoot mortality and shoot production, reacted differently to the two stress types we have simulated. Shoot death was accelerated by high temperatures, while the emergence of new shoots was slowed down by nutrient loading. Hence, both stressors are detrimental to the seagrass development, but operate through different mechanisms. The environment in SB is considered to be less disturbed than the environment in TY. However, Mejia et al. [19] measured higher nutrient concentrations in SB water and pore water than in TY water. It is therefore challenging to determine whether environmental conditions have driven TY seagrass to higher resilience, SB seagrass to higher fragility, or if phenological differences between the sites were at all shaped by environmental factors. Indeed, our experiment does not allow us to distinguish among several possible mechanisms such as selection, plasticity, gene flow or drift, which may have been responsible for the phenological differences between the two seagrass populations.

Our results also reveal a complex effect of temperature, nutrient concentrations and time on the epiphytic microbial community associated with *H. stipulacea*. Only 20% of the total compositional variance was attributed to factors which were accounted for or to interactions among them. Possibly, additional important independent factors, which were not accounted for, might have explained the remaining variance. For example, variability in leaf exudates within a seagrass population would likely be reflected in the microbial composition, and sediment microbiome dynamics may have directly modulated the nutrient measurements. Another possibility is that biotic interactions within the epiphytic communities were significant sources of variance. Depending on the initial epiphytic community compositions, such interactions would have had different consequences under similar conditions, particularly when taking the effect of ecological drift into account [52]. Under strong ecological drift, the resolution of abiotic perturbations in a microbial community is partially stochastic, especially if different taxa share functions and have equivalent fitnesses. The small size of the mesocosm, in comparison to the size of natural meadows and their environment, could have contributed to an increased ecological drift effect.

With the exception of time, the temperature had the largest effect on the bacterial composition, followed by the effects of the nutrient concentrations. The source site had a small and borderline significant effect. However, the unifrac distance distributions within and among source sites revealed that the TY communities, initially more conserved than the SB communities, diverged to a larger extent from one another than the SB epiphytic microbial communities, by the end of the experiment. This result coincided with lower mortality rates and higher growth rates in the TY seagrass and should thus be evaluated as a possible source of beneficial holobiont level plasticity of the TY seagrass in future studies. Still, the source site of the seagrass had a minor effect on the microbial community composition, in contrast with its effect on seagrass phenology, and the microbiota were mostly shaped by environmental factors. These results strongly support the “environmental fingerprint” hypothesis formulated by Mejia et al. [19], Rotini et al. [7] and Conte et al. [20] and highlight the early warning information that can be gained regarding exposure to stress, from monitoring the microbiota in wild meadows.

In terms of alpha diversity, only the combination of control temperatures (27°C) with enriched nutrients (the CTN treatment) caused a reduction in Faith PD, in comparison with the control treatment. The reduction in alpha diversity was explained mainly by the increase in Rhodobacterales and particularly by ASV 71a746. Based on a phylogenetic analysis, ASV 71a746 is most closely related to genbank accession AM691091, an extremophile from the East German Creek system in Canada [42], within *Roseobacter*, a genus normally containing temperate and polar species [43], and was first described in seaweed [44]. It is therefore possible that this ASV represents the southern edge of its congeneric distribution, can only flourish under the baseline temperature of 27°C, but effectively utilizes the excess nutrients. Interestingly, *Roseobacter* spp. reduce NO_3_^−^ to NO_2_^−^ [44], which is consistent with the enriched NO_2_^−^ measurements under the CTN scenario. *Granulosicoccus* ASV 9d7587, another cold water nitrate reducer, followed a similar pattern to that of the Rhodobacterales ASV 71a746. Concomitantly, Microtrchales, and Microtrichaceae in particular, also prevailed particularly under the CTN scenario. Microtrichaceae were found to be highly abundant in a partial nitrification - anammox system, where partial nitrification, such as that carried out by *Roseobacter*, produces NO_2_^−^, which is then used by Microtrichaceae to oxidize NH_4_^+^ [45]. Therefore, the dynamics of these microorganisms may serve as a buffer allowing *H. stipulacea* to cope with nitrogen enriched environments. Interestingly, the highest seagrass mortality was observed under high temperature treatments, and almost never under nutrient enrichment alone (CTN). The presence of these bacteria on seagrass leaves, and their increase under high nutrient concentrations with baseline temperatures, may represent a resilience mechanism of *H. stipulacea* to nitrogen-enriched environments, possibly allowing them to outcompete other seagrass species in anthropogenically disturbed, or less oligotrophic areas than the Gulf of Aqaba. For example, it would be interesting to test whether similar dynamics are absent from *Cymodocea nodosa*, which is rapidly displaced by *H. stipulacea* in the Mediterranean Sea, particularly in a disturbed harbour on the Tunisian coast [24]. Previous studies have reported that Red Sea *H. stipulacea* grew faster, forming denser meadows in polluted areas [29, 53], which is consistent with the low effect of nutrient loading on mortality that we report here. Such a buffering mechanism, however, would be sensitive to the increase in seawater temperatures, according to our results.

The experiment also sheds light on the response of cyanobacterial biofilms to each of the stressors. The relative abundance of Cyanobacteria increased by T8 under all treatments, probably due to the accumulation of nutrients in the aquaria, particularly under TN, due to the nutrient enrichment - even weeks after active loading ceased. They bloomed earliest under the CTN treatment, where temperatures were kept at baseline (27°C) but nutrients were actively enriched. *Rivularia* spp., to which the early blooming ASV belonged, was the most abundant cyanobacterial ASV. *Rivularia* spp. was already shown to be sensitive to nutrient loading [54], and accordingly this ASV was displaced by an ASV belonging to *Pseudanabaena* sp., which emerged at T8 under all treatments. A *Lyngbia* sp. ASV which also emerged at T8 under TN is known to form biofilms on seagrass leaves and reduce light availability [40]. Interestingly, the relative abundance of Cyanobacteria was a major contributor to the difference between TY and SB *in situ* [19]. With higher abundances in the SB meadow, where nutrient concentrations were higher. Lastly, cyanobacteria, although considered to either increase nitrogen assimilation [35] and limit light availability [40], their increased abundance under CTN does not seem to be detrimental to the seagrass in the mesocosm. However, light was not a limiting factor in our experiment, and cyanobacteria fix nitrogen only when it is not otherwise available [55] so in fact, they may participate in the buffering of nutrient loading.

## Conclusion

In this study, we were able to tease apart the impacts of environment dependent and host dependent factors on phenological and microbial properties of *H. stipulacea*, illustrating that the measured phenological properties are mostly host dependent while the epiphytic microbial composition is mostly environment dependent. This result supports the “environmental fingerprint” hypothesis raised in recent studies, and highlights the utility of microbiome shifts as bioindicators of nutrient exposure changes in *H. stipulacea* meadows, a fundamentally important component of the ecosystem in the Gulf of Aqaba. We further propose that bacterial community dynamics contribute to the holobiont’s plasticity by buffering nutrient effects when high concentrations are encountered, which may facilitate the range extension of *H. stipulacea* into new habitats, including northerner latitudes with less oligotrophic waters than in their native range. Our experiment demonstrated that when exposed to either stressor, plants from the TY population, a site with medium anthropogenic impacts, performed better than plants from the SB population, a less disturbed site, although understanding the process, which caused this differentiation, is beyond the scope of our experiment. Our findings may have important implications concerning the future of *H. stipulacea* as global climate changes progress and the importance of the holobiont perspective in understanding them.

## Methods

### Plant collection

Intact and healthy *H. stipulacea* plants bearing 5-6 shoots were collected in July 2019 from 6-8 m depth (Irradiances of 250 μmol photons m^−2^ S^−1^) by scuba diving. Two populations in the northern Gulf of Aqaba (Eilat, Israel) were visited: South Beach (SB; 29.497664°N 34.912737°E) and Tur-Yam Beach (TY; 29.516527°N 34.927205°E). At each site, 140 plants were collected from every 5-10 m to avoid pseudoreplicates. The plants were put in ziplock bags filled with seawater and were transported in a cooler box to the seagrass mesocosm facility at the Dead Sea Arava Science Center, where they were immersed in freshwater for 2-3 min and wiped off to remove organisms as much as possible with minimal damage to the plants. The plants were then inoculated by submerging them in natural seawater collected from TY. Six additional *H. stipulacea* were collected from each meadow on Dec 15 2019 and stored at −80°C until further processing to compare the post-acclimation mesocosm epiphytic communities (see below) with epiphytic communities in the field.

### Experimental setup

A mesocosm aquaria system (Fig. 1) was set up to simulate heatwave and eutrophication conditions, alongside control conditions. The system included four temperature baths, each controlled with a ProfiLux aquarium controller (GHL aquarium computers, Germany). Each bath contained five 60 L aquaria, layered with 6 cm of sieved and autoclaved natural coastal sediment, and filled with artificial seawater at a 40 practical salinity units concentration of Red Sea salt (www.redseafish.com). In each aquarium we planted 16-18 plants from each population (TY and SB) side by side, divided by a barrier between the two populations. Plants were acclimated for four weeks to baseline mesocosm conditions (27°C, 250 μmol photons m^−2^ S^−1^ at the water surface, 12hr light/day), allowing them to recover from the potential stress inflicted during plant collection, transportation and transplantation. Seagrasses were then exposed to four different treatments in each of the four baths: *i*) control water temperature with no nutrient enrichment (CTCN), *ii*) control water temperature with nutrient enrichment (CTN),*iii*) increased water temperatures (31°C) with no nutrient enrichment (TCN), and *iv*) the combination of the two stressors - increased water temperatures (31°C) with nutrient enrichment (TN). To initiate these experimental conditions at the end of the 4-weeks acclimation period (T_0_), using aquaria heaters, the temperature in two baths was gradually increased (0.8°C/day) from 27°C to 31°C, ~4°C above the average summer temperatures, simulating the increased temperature of the Red Sea at the end of the century [12, 56, 57]. The rest of the aquaria were left at control water temperatures of 27°C. Similarly, nutrients were gradually loaded in the aquaria of two baths, simulating eutrophication, by adding crushed slow-release fertilizer pellets (Osmocote, 17:11:10 N:P:K) twice a week, until reaching a final nitrate concentration of 20 μm L-1. Once reaching the target stress conditions of 31°C and 100μm nitrate (T1), these conditions were sustained for additional 5 weeks (T1-T4; the stress phase), before gradually reducing the temperature back to 27°C at a 0.8°C/day rate, and nutrient enrichment was stopped (T4). Once the baseline temperature of 27°C was regained in all the baths (T5), plants were allowed to recover from the stress conditions for 3 additional weeks (T5-T8). Throughout the experiment, water exchanges were made weekly (~10% of seawater volume) and light intensity and salinity was kept constant. Water temperatures were logged automatically every hour with GHL PT-1000 electrodes (GHL, Aquarium Computer, Germany; 2/bath), and manually daily (WTW 340i, WTW, Germany). Water samples were taken (50 ml, filtered through a 0.22 micron syringe filter, kept frozen) for future nutrient analysis. Nutrients were measured to confirm the establishment of nutrient concentration differences between the high and low nutrient treatments. It should be noted that the measured nutrients reflect not only the administered quantities but also the accumulation of nutrients and interactions with biotic factors.

### Phenological seagrass population descriptors

Seagrass phenology was a dependent variable in this experiment and was accounted for by quantifying the production of new shoots and the death of shoots. At the end of each time point (T_0_ to T_8_) the number of newly produced shoots and number of dead shoots since the previous time point were counted across each bath. The effect of temperature and nutrient stress treatments, source seagrass population and time on seagrass phenology was tested with a type-3 factorial anova, as implemented in StatModels [58].

### Nutrient concentration measurements

Mesocosm seawater samples for nutrient analysis were collected with plastic syringes (~250 mL) above the *H. stipulacea* shoots of each aquarium (n = 4). Seawater samples were instantly filtered using sterile syringe filters (cellulose acetate; 0.45 μm pore size; LABSOLUTE®) into HDPE vials and frozen at −80 °C. We measured the concentrations of NO_2_^−^, NO_x_ (NO_3_^−^ and NO_2_^−^), NH_4_^+^, PO_4_^3−^. Nutrient analyses were performed spectrophotometrically with a TECAN plate reader (Infinite 200 Pro microplate reader; Switzerland) following Laskov et al. [59]. The detection limits were 0.08, 0.32, 0.7 and 0.022 μM for NO_2_^−^, NO_x_ (NO_3_^−^ and NO_2_^−^), NH_4_^+^, PO_4_^3−^, respectively. The coefficient of variation was always < 3.4%.

### Epiphytic community samples collection

Epiphytic community samples were collected from both the mesocosm aquaria and from the *H. stipulacea* meadows at TY and SB, to study the bacterial dynamics in the experiment and to evaluate the relevance of the experimental results to the microbiota of the natural *H. stipulacea* populations. In the mesocosm, two leaves from the third shoot of one SB and one TY plant were collected from four tanks per treatment (n=4) in each of the following time points: following acclimation at the beginning of the experiment (T0), during the induced stress period (T3), at the end of the induced stress period (T5) and following the three weeks recovery period (T8). At each of these time points, one water sample was collected from each treatment. Third shoot leaves from wild and mesocosm plants were placed in DNeasy PowerSoil C1 solution (Qiagen) and sonicated for 3 minutes at 1.2 kHz in a sonicator bath. The C1 solution containing the sheared epiphytes underwent immediate DNA extraction. Water samples were filtered onto mixed cellulose esters 0.22-μm-pore-size filters, which were stored at −80°C until further processing.

### 16S rRNA metabarcoding

DNA was extracted from the epiphyte C1 solution using the DNeasy PowerSoil kit (Qiagen), and from water microbiome filters using the PowerWater DNA extraction kit (Qiagen), following the manufacturer’s instructions. Metabarcoding libraries were prepared with a two step PCR protocol. For the first PCR reaction (PCR1), the V4 16S rRNA region was amplified, following [60] et al. [60], with the forward primer 515f 5’-tcg tcg gca gcg tca gat gtg tat aag aga cag GGT GCC AGC MGC CGC GGT AA-3’ and the reverse primer 806R 5’-gtc tcg tgg gct cgg aga tgt gta taa gag aca gGA CTA CHV GGG TWT CTA AT-3’, along with artificial overhang sequences (lowercase). In the second PCR reaction (PCR2), sample specific barcode sequences and Illumina flow cell adapters were attached, using the forward primer ‘5-AAT GAT ACG GCG ACC ACC GAG ATC TAC ACt cgt cgg cag cgt cag atg tgt ata aga gac ag-’3 and the reverse primer ‘5-CAA GCA GAA GAC GGC ATA CGA GAT XXX XXX XXg tct cgt ggg ctc gg-’3’, including Illumina adapters (uppercase), overhang complementary sequences (lowercase), and sample specific DNA barcodes (‘X’ sequence). The PCR reactions were carried out in triplicate, with the KAPA HiFi HotStart ReadyMix PCR Kit (KAPA Biosystems), in a volume of 25 μl, including 2 μl of DNA template and following the manufacturer’s instructions. PCR1 started with a denaturation step of 3 minutes at 95 °C, followed by 30 cycles of 20 seconds denaturation at 98°C, 15 seconds of annealing at 55°C and 7 seconds polymerization at 72°C. The reaction was finalized with another minute-long polymerization step. PCR2 was carried out in a volume of 25 μl as well, with 2 μl of the PCR1 product as DNA template. It started with a 3-minutes denaturation step at 95°C, followed by 8 cycles of 20 seconds denaturation at 98°C, 15 seconds of annealing at 55°C and 7 seconds polymerization at 72°C. PCR2 was also finalized with another 60-second polymerization step. Products of PCR1 and PCR2 were purified using AMPure XP PCR product cleanup and size selection kit (Beckman Coulter), following the manufacturer’s instructions, and normalised based on Quant-iT PicoGreen (Invitrogen) quantifications. The fragment size distribution in the pooled libraries was examined on a TapeStation 4200 (Agilent) and the libraries were sequenced on an iSeq-100 Illumina platform, producing 150 bp paired end reads. Sequence data was deposited in the National Center for Biotechnology Information (NCBI) data bank, under BioProject accession PRJNA750596.

### Amplicon sequence variants (ASVs) and taxonomy assignment

All the analyses carried out for this study are available as a Jupyter notebook in a github repository (GitHub: https://git.io/JBBeV, Zenodo DOI: 10.5281/zenodo.5217277), along with the sequence data, intermediate and output files. The bioinformatics analysis was carried out within the Qiime2 [61] framework. DADA2 [62] was used to trim PCR primers, quality-filter, error correct, dereplicate and merge the read pairs, and to remove chimeric sequences, to produce the ASVs. Throughout the text, specific ASVs are referred to by the first 6 to 8 characters of their MD5 digests, which correspond with the biom table headers and the sequence IDs in the ASV fasta file. For taxonomic assignment, a naive Bayes classifier was trained using taxonomically identified reference sequences from the Silva 138 SSU-rRNA database [63] for the V4 fragments. All ASVs that were identified as mitochondrial or chloroplast sequences were filtered out from the feature table along with ASVs that had less than 30 occurrences across the dataset. An ASV phylogenetic tree was built with MAFFT 7.3 [64] for sequence alignment, and FastTree 2.1 [65], with the default masking options of the q2-phylogeny Qiime2 plugin.

### Microbial diversity analyses

ASVs shared between the epiphytic and planktonic communities were excluded from the biodiversity and differential abundance analyses of epiphytes. Microbial diversity was estimated based on Faith’s phylogenetic diversity [Faith PD; 66] for alpha diversity and weighted and unweighted UniFrac distance [67] matrices for beta diversity, all of which consider the phylogenetic relationships among ASVs. Ordination of the beta-diversity pairwise distances was carried out with a principal coordinates analysis [PCoA; 41, 68]. The contribution to bacterial compositional variance by time, temperature, nutrient concentrations and the source seagrass population was computed with the redundancy analysis anova procedure [69], as implemented in Vegan 2.5 [70]. Their contribution to alpha and beta diversity was additionally tested with a factorial ANOVA using the q2-longitudinal plugin [71]. P-values were corrected for multiple testing using the Benjamini-Hochberg procedure [72]. Corrected p-values are referred to as q-values throughout the text. All microbial diversity analyses were carried out using rarified tables. ANOVA tests were similarly carried out to test for significant differences between “within source site” and “among source site” pairwise UniFrac distance distributions.

### Differentially abundant and explanatory taxa and ASVs

Two groups of order level taxa were considered when testing for differentially abundant orders, including the eight most relatively-abundant orders at each time point as well as low relative-abundance orders. The Kruskal Wallis test [73] was used to test for differential abundances among the highly abundant orders, at each time point separately, and additional differently abundant orders were identified with the Analysis of composition of microbiomes (ANOCM) procedure [74]. ANCOM was used to test for differential abundance at the family level as well, and BiPlot [41] analyses were used to highlight “important” ASVs, following the importance definition of Legendre and Legendre [41], which best explained the weighted and unweighted UniFrac pairwise distance matrices.

### Core ASVs

Core ASVs were defined as ASVs found in all the wild samples and in all the samples of at least one of the treatments at time point zero. Similar core communities were identified for SB and TY epiphyte samples separately. To ensure that the regarded core communities bear relevance to the field, ASVs that were not present in field samples from the SB and TY sites were excluded from the core microbiome.

## Ethics approval and consent to participate

Not applicable

## Consent for publication

Not applicable

## Availability of data and materials

The datasets generated and analysed during the current study are available in the National Center for Biotechnology Information (NCBI) BioProject repository under the accession number PRJNA750596. Data and script are archived as a GitHub release (https://git.io/JBBeV, DOI: 10.5281/zenodo.5217277).

## Authors’ contributions

GW and PBC designed the study. PBC, TAG, AS and GW collected the samples. PBC and GW carried out the nutrient measurements and temperature measurements. PBC and TAG carried out the phenological measurements. TY and RSA carried out the molecular lab work. AS and TY analysed the data. AS, GW, PBC and TY wrote the manuscript. All authors read and approved the final manuscript.

## Acknowledgements

We thank Israel Ministry of Science and Technology for their support of Dead Sea and Arava Science Center.

## Supplementary information

Figure S1: Alpha rarefaction curves of epiphytic microbial communities in samples of each mesocosm treatment.

Figure S2: Alpha and beta diversity of mesocosm epiphyte and water samples. Faith’s phylogenetic diversity distributions in water and epiphyte samples are presented as a box plot. Weighted Unifrac distance based PCoA and Biplot are presented as ordination of PC1 and PC2.

Figure S3: Diverging community compositions among treatments. Weighted UniFrac distance based principal coordinate analyses (PCoA) of endophyte samples from T0 (A), T3 (B), T5 (C) and T8 (D). CTCN - control temperatures (27°C) and control nutrients (no enrichment). CTN - control temperatures (27°C) with nutrient enrichment. TCN - heatwave (31°C) without nutrient enrichment. TN - heatwave with nutrient enrichment. Time points T0 and T8 had baseline temperatures and no active nutrient enrichment in all baths. The percent total variance accounted for by each coordinate is indicated on the corresponding axis. The most important ASVs, following the importance definition by Legendre and Legendre [41], are represented by BiPlot analyses (gray arrows) and their taxonomic identifications are noted in the legend.

Figure S4: Family level dynamics of ANCOM families. CTCN - control temperatures (27°C) and control nutrients (no enriching). CTN - control temperatures with nutrient enriching . TCN - heatwave (31°C) without nutrient enriching. TN - heatwave with nutrient enriching. Timepoints T0 and T8 had baseline temperatures and no active nutrient enrichment in all treatments.

Figure S5: A phylogenetic tree of Rhodobacterales ASVs (red) along with reference sequences from the SILVA database (black) and their isolation source (green). Black bullets at the base of nodes represent a bootstrap percentage or 70 or higher.

Figure S6: A phylogenetic tree of Saprospiraceae ASVs (red) along with reference sequences from the SILVA database (black) and their isolation source (green). Black bullets at the base of nodes represent a bootstrap percentage or 70 or higher.

Figure S7: A phylogenetic tree of Cyanobacteria ASVs (red) along with reference sequences from the SILVA database (black) and their isolation source (green). Black bullets at the base of nodes represent a bootstrap percentage or 70 or higher.

Figure S8: Dynamics of key features. Values are normalised by the peak relative abundance of each feature separately.

Figure S9: Relative abundance dynamics of core ASVs.

Figure S10: Relative abundance dynamics of TY core ASVs.

Figure S11: Relative abundance dynamics of SB core ASVs.

## References

1. Al-Rousan S, Al-Horani F, Eid E, Khalaf M. Assessment of seagrass communities along the Jordanian coast of the Gulf of Aqaba, Red Sea. Mar Biol Res. 2011;7:93–9.

2. Winters G, Edelist D, Shem-Tov R, Beer S, Rilov G. A low cost field-survey method for mapping seagrasses and their potential threats: an example from the northern Gulf of Aqaba, Red Sea. Aquat Conserv. 2017;27:324–39.

3. Sharon Y, Levitan O, Spungin D, Berman-Frank I, Beer S. Photoacclimation of the seagrass *Halophila stipulacea* to the dim irradiance at its 48-meter depth limit. Limnol Oceanogr. 2011;56:357–62.

4. Den-Hartog C. The Sea-Grasses of the World. Amsterdam: North-Holland; 1970.

5. Den-Hartog C, Kuo J. Taxonomy and biogeography of seagrasses. In: Larkum AWD, Orth RJ, Duarte CM, editors. Seagrasses: Biology, Ecology and Conservation. Dordrecht: Springer; 2006. p. 1–23.

6. Sharon Y, Silva J, Santos R, Runcie JW, Chernihovsky M, Beer S. Photosynthetic responses of *Halophila stipulacea* to a light gradient. II. Acclimations following transplantation. Aquat Biol. 2009;7:153–7.

7. Rotini A, Mejia AY, Costa R, Migliore L, Winters G. Ecophysiological plasticity and bacteriome shift in the seagrass *Halophila stipulacea* along a depth gradient in the northern Red Sea. Front Plant Sci. 2017;7.

8. Unsworth RKF, Collier CJ, Henderson GM, McKenzie LJ. Tropical seagrass meadows modify seawater carbon chemistry: implications for coral reefs impacted by ocean acidification. Environ Res Lett. 2012;7:024026.

9. Lamb JB, van de Water JAJM, Bourne DG, Altier C, Hein MY, Fiorenza EA, et al. Seagrass ecosystems reduce exposure to bacterial pathogens of humans, fishes, and invertebrates. Science. 2017;355:731–3.

10. Nordlund LM, Jackson EL, Nakaoka M, Samper-Villarreal J, Beca-Carretero P, Creed JC. Seagrass ecosystem services - What’s next? Mar Pollut Bull. 2018;134:145–51.

11. Liao E, Lu W, Yan X-H, Jiang Y, Kidwell A. The coastal ocean response to the global warming acceleration and hiatus. Sci Rep. 2015;5:16630.

12. Fine M, Gildor H, Genin A. A coral reef refuge in the Red Sea. Glob Chang Biol. 2013;19:3640–7.

13. Silverman J, Gildor H. The residence time of an active versus a passive tracer in the Gulf of Aqaba: A box model approach. Journal of Marine Systems. 2008;71:159–70. doi:10.1016/j.jmarsys.2007.06.007.

14. Waycott M, Duarte CM, Carruthers TJB, Orth RJ, Dennison WC, Olyarnik S, et al. Accelerating loss of seagrasses across the globe threatens coastal ecosystems. Proc Natl Acad Sci U S A. 2009;106:12377–81.

15. McKenzie LJ, Long L, Coles RG, Roder CA. Seagrass-Watch: Community based monitoring of seagrass resources. Biol Mar Mediterr. 2000;7:393–6.

16. Short FT, McKenzie LJ, Coles RG, Gaeckle JL. SeagrassNet manual for scientific monitoring of seagrass habitat – western pacific edition. 2004. https://scholars.unh.edu/jel/399/. Accessed 4 Aug 2021.

17. Short FT, McKenzie LJ, Coles RG, Gaeckle JL. SeagrassNet Manual for scientific monitoring of seagrass habitat--Caribbean Edition. Durham, NH: University of New Hampshire. 2005.

18. Macreadie PI, Schliep MT, Rasheed MA, Chartrand KM, Ralph PJ. Molecular indicators of chronic seagrass stress: A new era in the management of seagrass ecosystems? Ecol Indic. 2014;38:3.

19. Mejia AY, Rotini A, Lacasella F, Bookman R, Thaller MC, Shem-Tov R, et al. Assessing the ecological status of seagrasses using morphology, biochemical descriptors and microbial community analyses. A study in *Halophila stipulacea* (Forsk.) Aschers meadows in the northern Red Sea. Ecol Indic. 2016;60:1150–63.

20. Conte C, Rotini A, Manfra L, D’Andrea MM, Winters G, Migliore L. The seagrass holobiont: what we know and what we still need to disclose for its possible use as an ecological indicator. Water. 2021;13:406.

21. Lipkin Y. *Halophila stipulacea*, a review of a successful immigration. Aquat Bot. 1975;1:203–15.

22. Lipkin Y. Quantitative aspects of seagrass communities, particularly of those dominated by *Halophila stipulacea*, in Sinai (Northern Red Sea). Aquat Bot. 1979;7:119–28.

23. Gambi MC, Barbieri F, Bianchi CN. New record of the alien seagrass *Halophila stipulacea* (Hydrocharitaceae) in the western Mediterranean: a further clue to changing Mediterranean Sea biogeography. Mar Biodivers Rec. 2009;2.

24. Sghaier YR, Zakhama-Sraieb R, Benamer I, Charfi-Cheikhrouha F. Occurrence of the seagrass *Halophila stipulacea* (Hydrocharitaceae) in the southern Mediterranean Sea. Bot Mar. 2011;54:575–82.

25. Beca-Carretero P, Teichberg M, Winters G, Procaccini G, Reuter H. Projected rapid habitat expansion of tropical seagrass species in the Mediterranean Sea as climate Change Progresses. Front Plant Sci. 2020;11:555376.

26. Willette DA, Chalifour J, Debrot AOD, Engel MS, Miller J, Oxenford HA, et al. Continued expansion of the trans-Atlantic invasive marine angiosperm *Halophila stipulacea* in the Eastern Caribbean. Aquat Bot. 2014;112:98–102.

27. Steiner SCC, Willette DA. The expansion of *Halophila stipulacea* (Hydrocharitaceae,Angiospermae) is changing the seagrass landscape in the commonwealth of Dominica, Lesser Antilles. Caribb Nat. 2015;22:1–19.

28. Beca-Carretero P, Guihéneuf F, Winters G, Stengel DB. Depth-induced adjustment of fatty acid and pigment composition suggests high biochemical plasticity in the tropical seagrass Halophila stipulacea. Mar Ecol Prog Ser. 2019;608:105–17.

29. Beca-Carretero P, Rotini A, Mejia A, Migliore L, Vizzini S, Winters G. *Halophila stipulacea* descriptors in the native area (Red Sea): A baseline for future comparisons with native and non-native populations. Mar Environ Res. 2020;:104828.

30. Winters G, Beer S, Willette DA, Viana IG, Chiquillo KL, Beca-Carretero P, et al. The Tropical seagrass *Halophila stipulacea*: reviewing what we know from its native and invasive habitats,alongside identifying knowledge gaps. Front Mar Sci. 2020;7:300.

31. Rosenberg E, Zilber-Rosenberg I. Microbes drive evolution of animals and plants: the hologenome concept. MBio. 2016;7:e01395.

32. Castell W zu, Fleischmann F, Heger T, Matyssek R. Shaping theoretic foundations of holobiont-like systems. In: Lüttge U, Cánovas FM, Matyssek R, editors. Progress in Botany 77. Springer International Publishing; 2016. p. 219–44.

33. Roughgarden J, Gilbert SF, Rosenberg E, Zilber-Rosenberg I, Lloyd EA. Holobionts as units of selection and a model of their population dynamics and evolution. Biol Theory. 2018;13:44–65.

34. Ugarelli K, Chakrabarti S, Laas P, Stingl U. The seagrass holobiont and its microbiome. Microorganisms. 2017;5.

35. Tarquinio F, Bourgoure J, Koenders A, Laverock B, Säwström C, Hyndes GA. Microorganisms facilitate uptake of dissolved organic nitrogen by seagrass leaves. ISME J. 2018;12:2796–800.

36. Welsh DT. Nitrogen fixation in seagrass meadows: Regulation, plant-bacteria interactions and significance to primary productivity. Ecology Letters. 2000;3:58–71. doi:10.1046/j.1461-0248.2000.00111.x.

37. Brodersen KE, Siboni N, Nielsen DA, Pernice M, Ralph PJ, Seymour J, et al. Seagrass rhizosphere microenvironment alters plant-associated microbial community composition. Environ Microbiol. 2018;20:2854–64.

38. Pereg LL, Lipkin Y, Sar N. Different niches of the *Halophila stipulacea* seagrass bed harbor distinct populations of nitrogen fixing bacteria. Mar Biol. 1994.

39. Uku J, Björk M, Bergman B, Díez B. Characterization and comparison of prokaryotic epiphytes associated with three east African seagrasses. J Phycol. 2007;43:768–79.

40. Tiling K, Proffitt CE. Effects of *Lyngbya majuscula* blooms on the seagrass *Halodule wrightii* and resident invertebrates. Harmful Algae. 2017;62:104–12.

41. Legendre P, Legendre L. Numerical Ecology. third. London: Elsevier; 2012.

42. Csotonyi JT, Swiderski J, Stackebrandt E, Yurkov VV. Novel halophilic aerobic anoxygenic phototrophs from a Canadian hypersaline spring system. Extremophiles. 2008;12:529–39.

43. Buchan A, González JM, Moran MA. Overview of the marine *Roseobacter* lineage. Appl Environ Microbiol. 2005;71:5665–77.

44. Shiba T. *Roseobacter litoralis* gen. nov., sp. nov., and *Roseobacter denitrificans* sp. nov., aerobic pink-pigmented bacteria which contain bacteriochlorophyll A. Syst Appl Microbiol. 1991;14:140–5.

45. Wang B, Wang Z, Wang S, Qiao X, Gong X, Gong Q, et al. Recovering partial nitritation in a PN/A system during mainstream wastewater treatment by reviving AOB activity after thoroughly inhibiting AOB and NOB with free nitrous acid. Environ Int. 2020;139:105684.

46. Walker AM, Leigh MB, Mincks SL. Patterns in benthic microbial community structure across environmental gradients in the beaufort sea shelf and slope. Front Microbiol. 2021;12:581124.

47. Konstantinou D, Voultsiadou E, Panteris E, Gkelis S. Revealing new sponge-associated cyanobacterial diversity: Novel genera and species. Mol Phylogenet Evol. 2021;155:106991.

48. Sekar R, Mills DK, Remily ER, Voss JD, Richardson LL. Microbial communities in the surface mucopolysaccharide layer and the black band microbial mat of black band-diseased Siderastrea siderea. Appl Environ Microbiol. 2006;72:5963–73.

49. Harris JK, Caporaso JG, Walker JJ, Spear JR, Gold NJ, Robertson CE, et al. Phylogenetic stratigraphy in the Guerrero Negro hypersaline microbial mat. ISME J. 2013;7:50–60.

50. Lee K, Lee HK, Choi T-H, Kim K-M, Cho J-C. Granulosicoccaceae fam. nov., to include *Granulosicoccus antarcticus* gen. nov., sp. nov., a non-phototrophic, obligately aerobic chemoheterotroph in the order Chromatiales, isolated from Antarctic seawater. J Microbiol Biotechnol. 2007;17:1483–90.

51. Leiva-Dueñas C, Martínez Cortizas A, Piñeiro-Juncal N, Díaz-Almela E, Garcia-Orellana J, Mateo MA. Long-term dynamics of production in western Mediterranean seagrass meadows: Trade-offs and legacies of past disturbances. Sci Total Environ. 2021;754:142117.

52. Gilbert B, Levine JM. Ecological drift and the distribution of species diversity. Proc Biol Sci. 2017;284. doi:10.1098/rspb.2017.0507.

53. Azcárate-García T, Beca-Carretero P, Villamayor B, Stengel DB, Winters G. Responses of the seagrass *Halophila stipulacea* to depth and spatial gradients in its native region (Red Sea): Morphology, in situ growth and biomass production. Aquat Bot. 2020;165:103252.

54. Uku J, Björk M. The distribution of epiphytic algae on three Kenyan seagrass species. S Afr J Bot. 2001;67:475–82.

55. O’Neil JM, Davis TW, Burford MA, Gobler CJ. The rise of harmful cyanobacteria blooms: The potential roles of eutrophication and climate change. Harmful Algae. 2012;14:313–34.

56. Nguyen HM, Savva I, Kleitou P, Kletou D, Lima FP, Sapir Y, et al. Seasonal dynamics of native and invasive *Halophila stipulacea* populations—A case study from the northern Gulf of Aqaba and the eastern Mediterranean Sea. Aquat Bot. 2020;162:103205.

57. Nguyen HM, Yadav NS, Barak S, Lima FP, Sapir Y, Winters G. Responses of invasive and native populations of the seagrass *Halophila stipulacea* to simulated climate change. Front Mar Sci. 2020;6:812.

58. Seabold S, Perktold J. Statsmodels: Econometric and statistical modeling with python. In: 9th Python in Science Conference. 2010.

59. Laskov C, Herzog C, Lewandowski J, Hupfer M. Miniaturized photometrical methods for the rapid analysis of phosphate, ammonium, ferrous iron, and sulfate in pore water of freshwater sediments. Limnol Oceanogr Methods. 2007;5:63–71.

60. Toju H, Kurokawa H, Kenta T. Factors influencing leaf- and root-associated communities of bacteria and fungi across 33 plant orders in a grassland. Front Microbiol. 2019;10:241.

61. Bolyen E, Rideout JR, Dillon MR, Bokulich NA, Abnet CC, Al-Ghalith GA, et al. Reproducible,interactive, scalable and extensible microbiome data science using QIIME 2. Nat Biotechnol. 2019;37:852–7.

62. Callahan BJ, McMurdie PJ, Rosen MJ, Han AW, Johnson AJA, Holmes SP. DADA2: High-resolution sample inference from Illumina amplicon data. Nat Methods. 2016;13:581–3.

63. Quast C, Pruesse E, Yilmaz P, Gerken J, Schweer T, Yarza P, et al. The SILVA ribosomal RNA gene database project: improved data processing and web-based tools. Nucleic Acids Res. 2013;41 Database issue:D590–6.

64. Katoh K, Standley DM. MAFFT multiple sequence alignment software version 7: improvements in performance and usability. Mol Biol Evol. 2013;30:772–80.

65. Price MN, Dehal PS, Arkin AP. FastTree 2-approximately maximum-likelihood trees for large alignments. PLoS One. 2010;5:e9490.

66. Faith DP. Conservation evaluation and phylogenetic diversity. Biol Conserv. 1992;61:1–10.

67. Lozupone C, Knight R. UniFrac: a new phylogenetic method for comparing microbial communities. Appl Environ Microbiol. 2005;71:8228–35.

68. Halko N, Martinsson P, Shkolnisky Y, Tygert M. An algorithm for the principal component analysis of large data sets. SIAM J Sci Comput. 2011;33:2580–94.

69. McCune B. Influence of noisy environmental data on canonical correspondence analysis. Ecology. 1997;78:2617–23.

70. Oksanen J, Blanchet FG, Friendly M, Kindt R, Legendre P, McGlinn D, et al. vegan: Community Ecology Package. 2020.

71. Bokulich NA, Zhang Y, Dillon M, Rideout JR, Bolyen E, Li H, et al. q2-longitudinal: a QIIME 2 plugin for longitudinal and paired-sample analyses of microbiome data. Cold Spring Harbor Laboratory. 2017;:223974.

72. Benjamini Y, Hochberg Y. Controlling the false discovery rate: a practical and powerful approach to multiple testing. J R Stat Soc B. 1995;57:289–300.

73. Kruskal WH, Wallis AW. Use of ranks in one-criterion variance analysis. J Am Stat Assoc. 1952;47:583–621.

74. Mandal S, Van Treuren W, White RA, Eggesbø M, Knight R, Peddada SD. Analysis of composition of microbiomes: a novel method for studying microbial composition. Microb Ecol Health Dis. 2015;26:27663.

