## Supplementary figures and images for "Teasing apart the host-related, nutrient-related and temperature-related effects shaping the phenology and microbiome of the tropical seagrass *Halophila stipulacea*"

### Figure S1

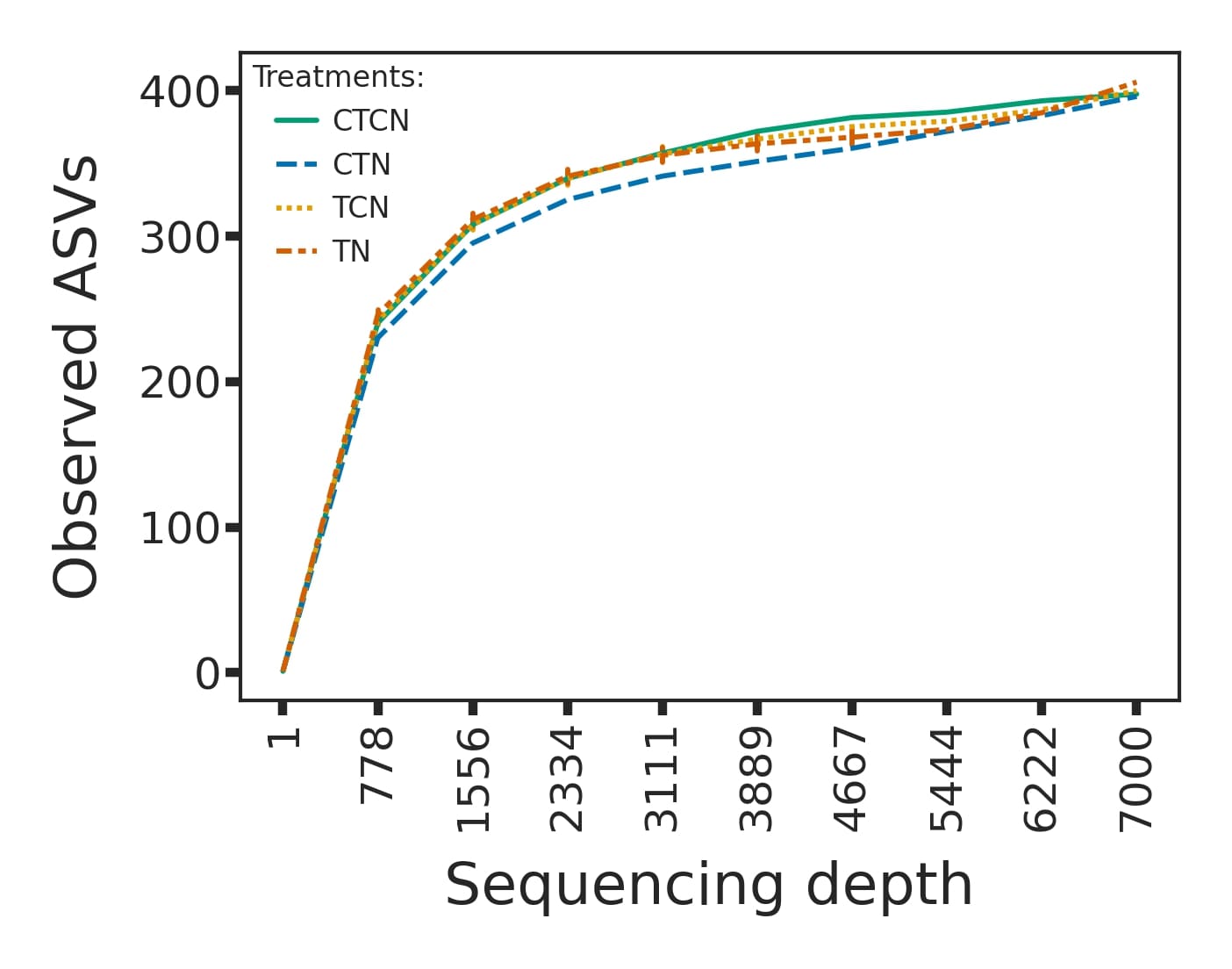

### Figure S2

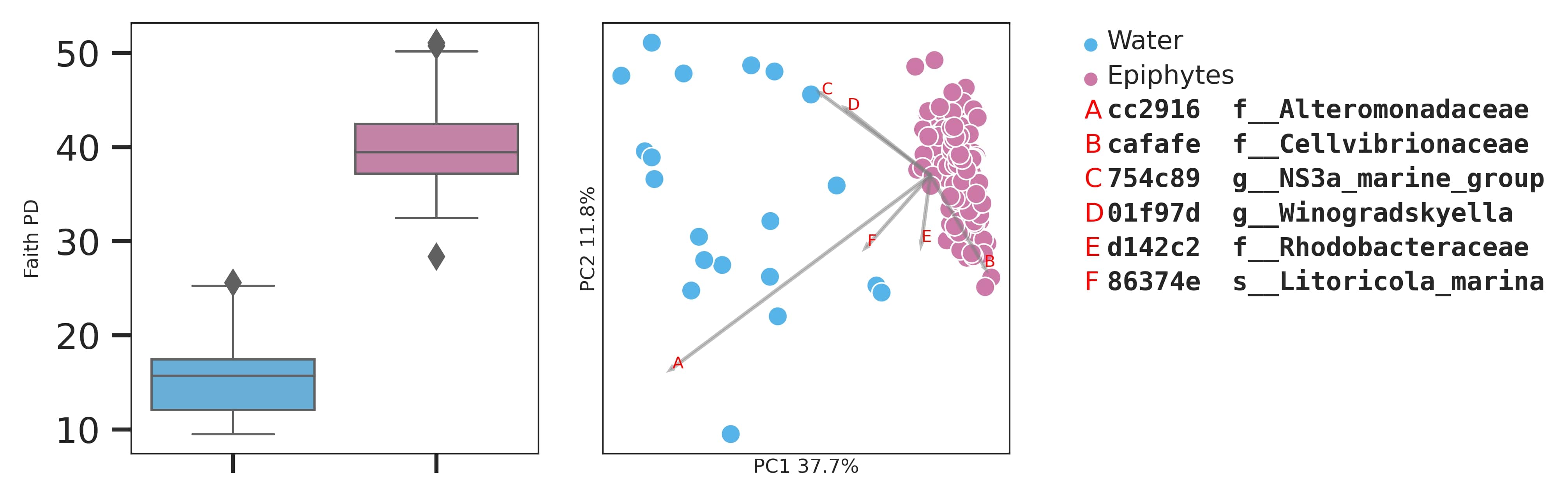

### Figure S3

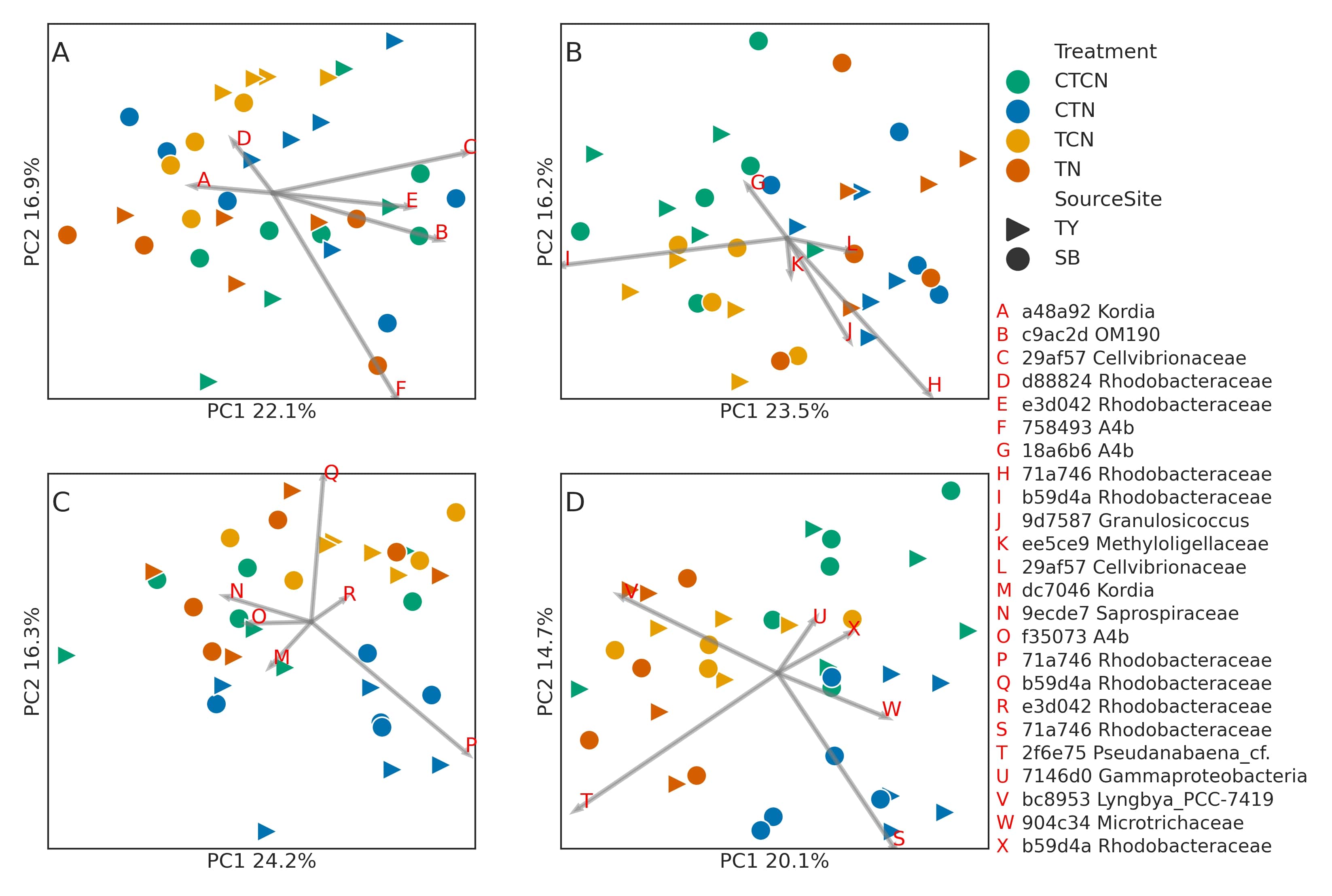

### Figure S4

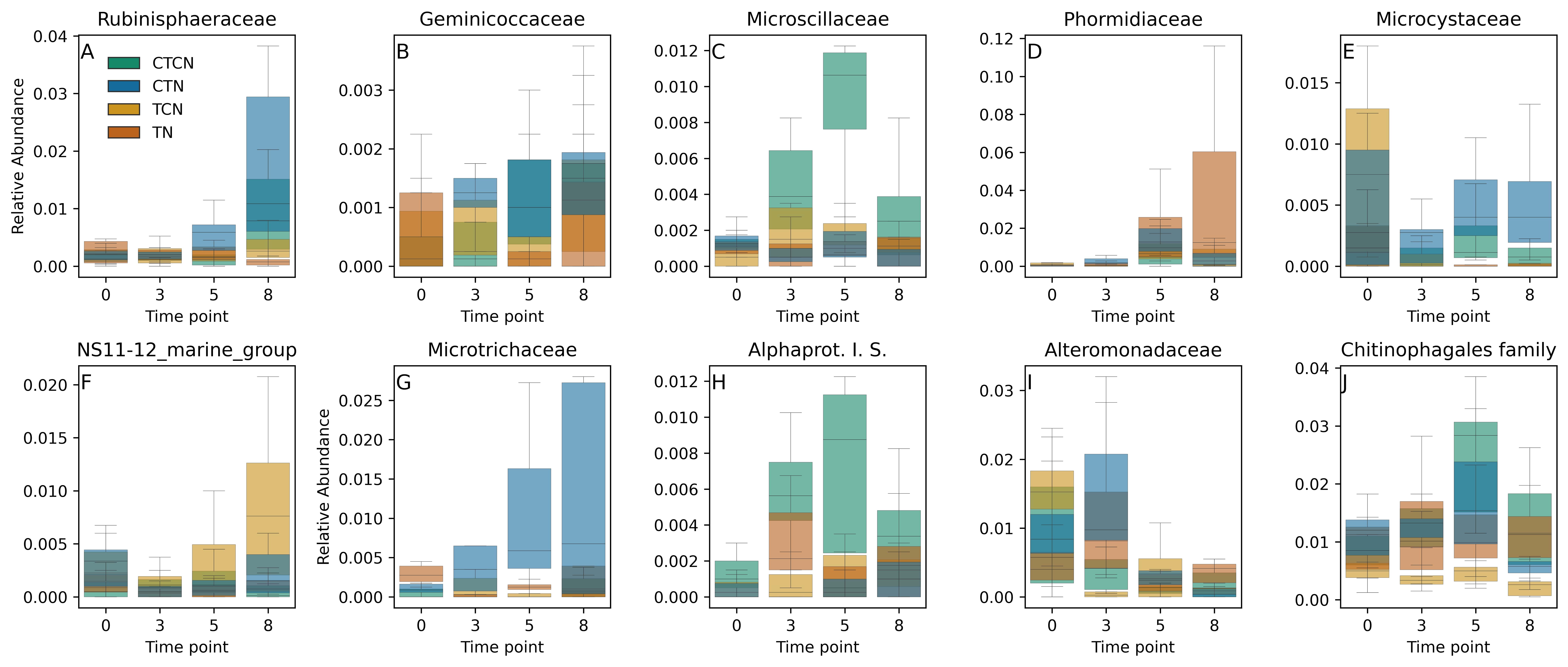

### Figure S5

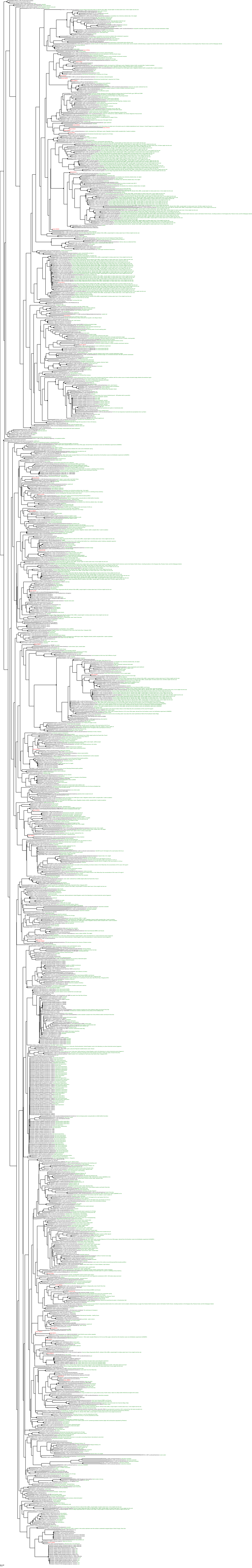

### Figure S6

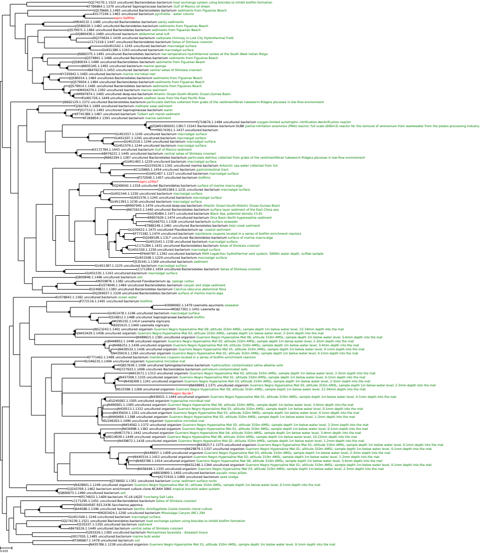

### Figure S7

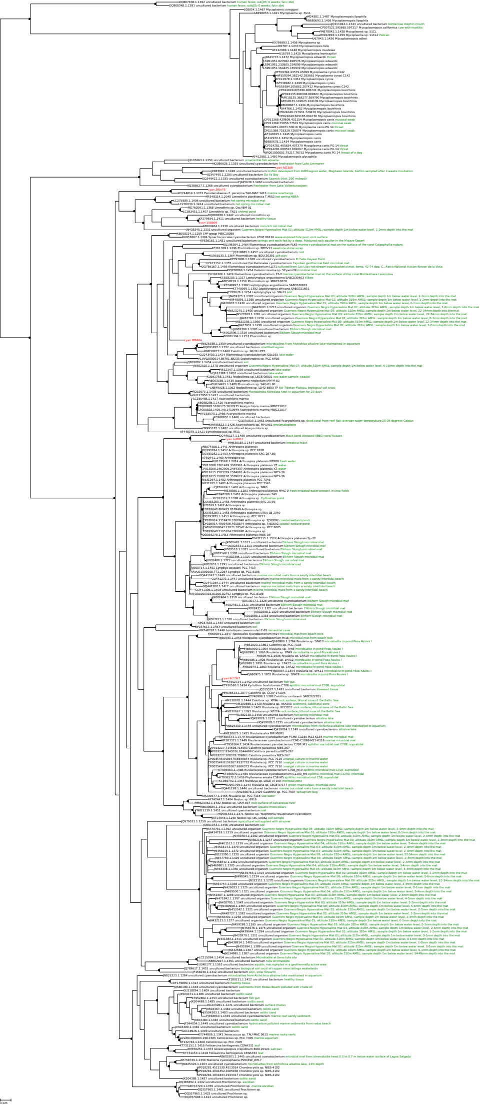

### Figure S8

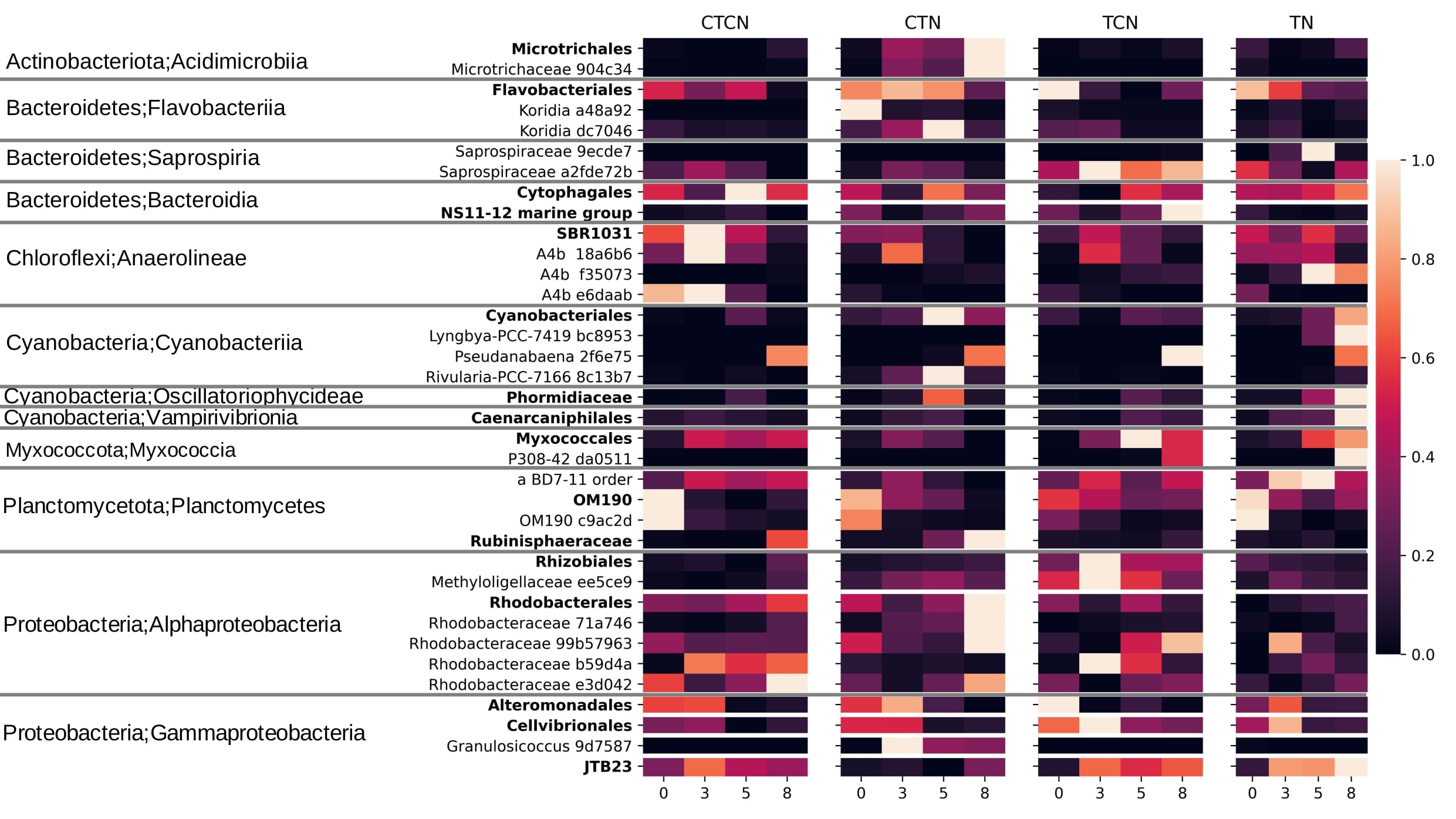

### Figure S9

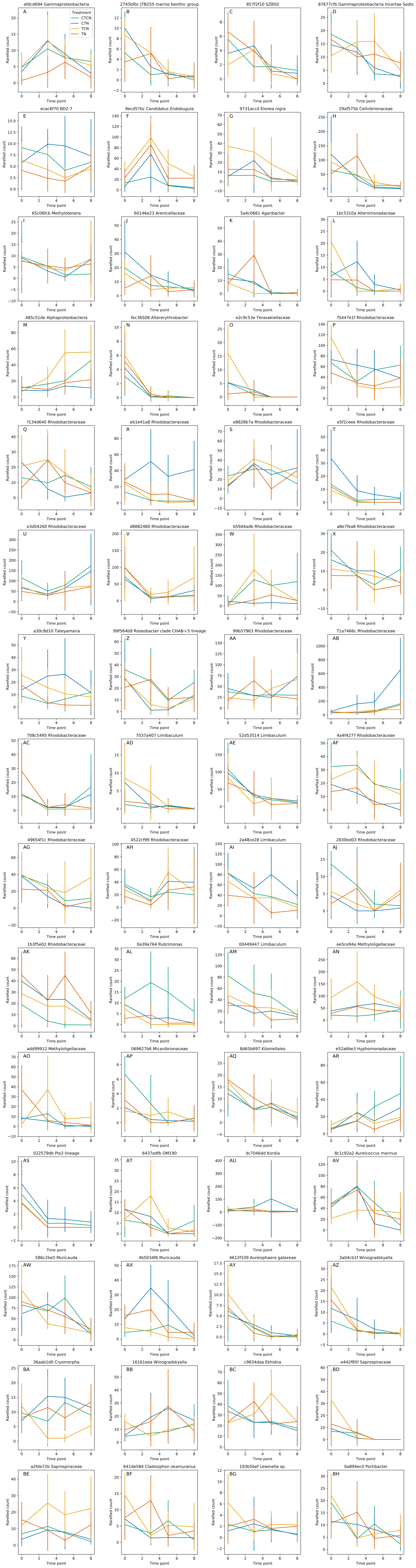

### Figure S10

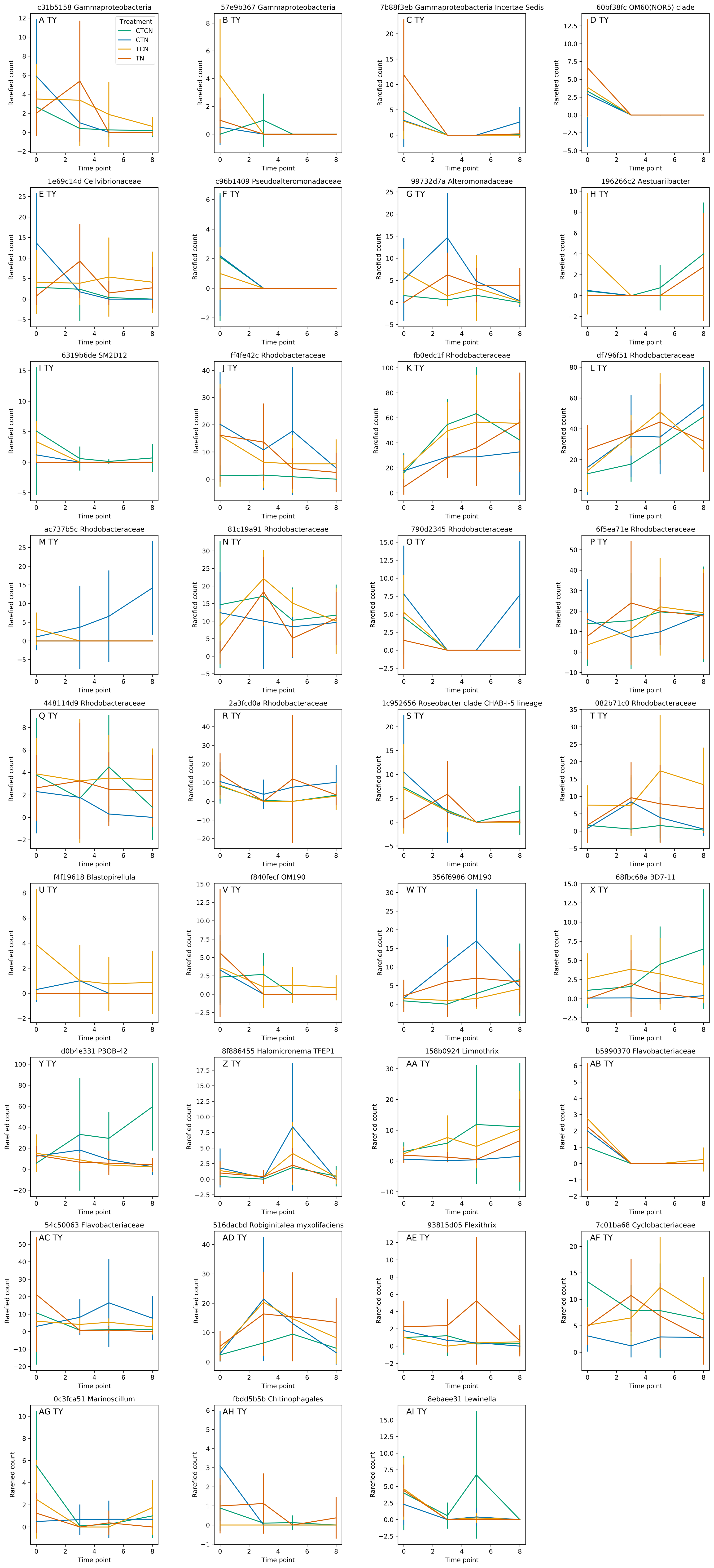

### Figure S11

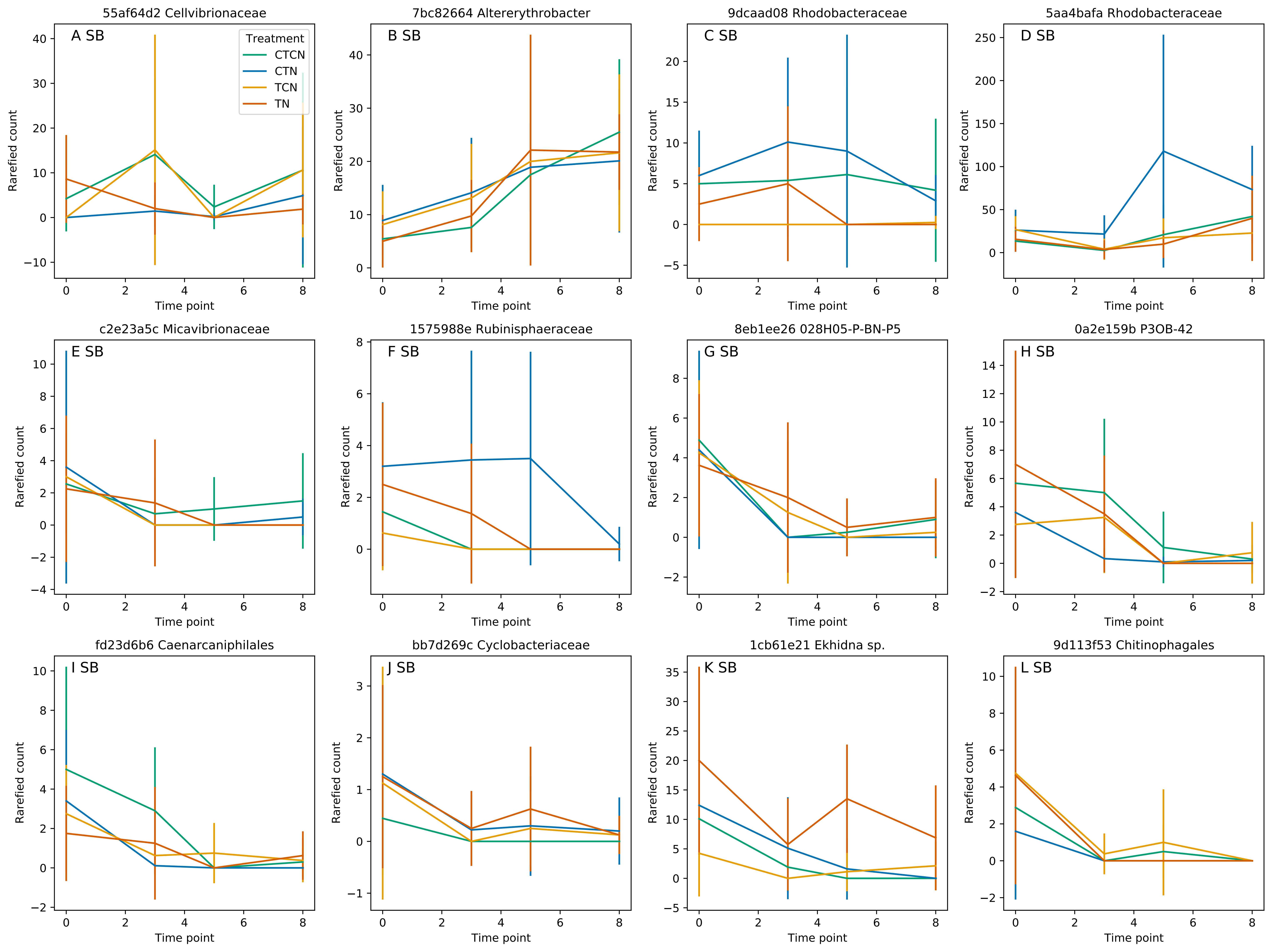
